# APOBEC3G Splicing Defects in Nonhuman Primate Models Result in Disparate Viral Mutational Profiles Relative to Humans

**DOI:** 10.64898/2026.08.06.743345

**Authors:** Armando D. Mendez, Rachael Springman-Rodriguez, Ayub Bokani, Fiona Carter-Tod, Niloofar Haghjoo, Margarita Rzhetskaya, Colin Rorex, Jake D. Lehle, Mohadeseh Soleimanpour, Luis Ferrández-Peral, Hanjing Yang, Michael A. Carpenter, Rajesh Thippeshappa, Sebla Kutluay, Rick N. McLaughlin, Mahesh Mohan, Binhua Ling, Luis D. Giavedoni, Maria Rodriguez-Barradas, Reuben S. Harris, Xiaojiang Chen, Susan T. Weintraub, Judd F. Hultquist, Diako Ebrahimi

## Abstract

Nonhuman primates (NHPs), particularly macaques, are indispensable models for studying human infectious diseases due to their close immunological and physiological similarities. Understanding species-specific molecular differences is essential for maximizing the translational value of these models. Here we report that *APOBEC3G (A3G)*, a potent antiviral restriction factor and the major source of genetic variations in HIV, exhibits a widespread mRNA splicing defect in the Cercopithecinae subfamily, which includes the commonly used NHP models. Driven by intronic polymorphisms, this splicing defect substantially reduces A3G protein levels and consequently results in a markedly reduced A3G-mediated mutation signatures, fewer defective viral genomes, and greater viral diversification in SIV compared to HIV. This species-specific effect is not restricted to lentiviruses: reduced A3G signatures have also been reported in simian foamy virus and simian T-cell leukemia virus, suggesting broader effects across primate retroviruses. These findings reveal a lineage-specific alteration in a major antiviral restriction factor, with important implications for viral restriction, evolution, drug resistance, and immune evasion. They also highlight the importance of incorporating naturally occurring genetic variation into NHP model selection to improve the reproducibility, translational fidelity, and biological relevance of preclinical research.

## INTRODUCTION

Nonhuman primates (NHPs) - particularly members of the Cercopithecinae subfamily of Old World monkeys (OWMs) such as rhesus macaques, pig-tailed macaques, and cynomolgus macaques, are indispensable animal models for studying a wide range of human infectious diseases and their causative agents, including poliomyelitis (poliovirus),^1^ measles (measles virus),^2^ Ebola virus disease (Ebola virus),^3,4^ human immunodeficiency virus/acquired immunodeficiency syndrome (HIV/AIDS; HIV-1),^5–13^ influenza (influenza viruses),^14^ Zika virus infection (Zika virus),^15,16^ malaria (*Plasmodium* spp.),^17^ and, more recently, coronavirus disease 2019 (COVID-19; SARS-CoV-2).^18,19^ The original selection of these primates as human disease models was primarily based on their physiological and immunological similarities to humans, ease of access and breeding, and manageable body size. Rhesus macaques have become a primary model for HIV research as SIV-infected macaques show CD4⁺ T-cell depletion, chronic immune activation, and latent reservoirs that are also seen in HIV infection.^20,21^

Although humans and NHPs share broad physiological and immunological similarities, fundamental molecular differences that influence immune responses, disease pathogenesis, and the translational relevance of these models have been documented.^22,23^ For instance, rhesus and cynomolgus TRIM5α isoforms can bind to the HIV capsid and mediate viral restriction, but this mechanism is absent in humans due to a couple of key amino acid substitutions that result in significant disease progression differences.^24–26^

A recent study reported an mRNA splicing defect in *APOBEC3G* (A3G) of Northern pig-tailed macaques.^27^ APOBEC3 (A3) enzymes constitute a vital component of the innate antiviral immune response, comprising seven family members (A3A/B/C/D/F/G/H) with broad antiviral activities mediated through cytosine deamination of viral genomes as well as through deamination-independent mechanisms.^28–33^ These enzymes act as critical barriers to cross-species transmission of lentiviruses, as evidenced by their evolutionary arms race with lentiviral Vif proteins, which target A3 enzymes for ubiquitin-mediated degradation.^34–36^ A3G is the most potent HIV restriction factor and the primary source of HIV mutations.^37,38^ This enzyme preferentially induces G-to-A mutations in a GG dinucleotide context (GG-to-AG/AA), often introducing premature termination codons (PTCs) that severely compromise viral fitness.^39–44^ By contrast, other A3 enzymes favor mutations within GA dinucleotides (GA-to-AA), which is much less likely to generate PTCs.^41–43,45–47^ At low frequency, A3-induced mutations can contribute to viral diversification, drug resistance, and immune evasion.^48–50^

The critical importance of A3G as a broad antiviral protein, a major lentiviral restriction factor, and a key source of viral genetic diversity promoted us to investigate the breadth and implications of the A3G splicing defect across diverse NHP species. Using a multi-omics approach, we found that the *A3G* mRNA splicing defect is highly prevalent and specific to the Cercopithecinae subfamily of Old World monkeys (OWMs), which comprises the species most widely used as human disease models.^51^ This defect results from aberrant exonization of an intronic region containing two PTCs, effectively abolishing A3G protein production as verified by mass spectrometry and ribosomal profiling. Consequently, unlike HIV in humans, SIV mutation spectra in these NHPs are dominated by GA-to-AA signatures from other A3 enzymes rather than the GG-to-AG mutations characteristic of A3G activity. This shift results in fewer defective viruses and greater viral diversification in SIV compared to HIV. These findings have implications beyond HIV and SIV. Similar patterns have been observed in other viruses: monkey foamy viruses and simian T-cell leukemia virus type 1 (STLV-1) show fewer A3G-associated GG-to-AG/AA mutations than their human counterparts, zoonotic foamy viruses and human T-cell leukemia virus type 1 (HTLV-1).^52–54^ Taken together, these findings highlight the importance of incorporating species-specific A3 variation in NHP models with implications for studies of viral diversification, drug resistance, immune evasion, vaccine efficacy, and therapeutic development.

## METHODS

### *Hypermutation profiling of HIV and SIV sequences from LANL databases* HIV sequences

We downloaded 932,134 HIV-1 sequences from the Los Alamos National Laboratories (LANL) HIV databases (accessed 12 Jan 2024). Sequences lacking donor IDs were excluded. Donors with fewer than three non-hypermutated (normal) sequences or no hypermutated sequences were discarded. This yielded 636,916 sequences from 588 donors, including 6,152 sequences annotated as hypermutated. For each hypermutated HIV sequence from each donor, we constructed a consensus sequence using 3-5 non-hypermutated sequences that shared ≥60% genomic overlap with the hypermutated sequence and exhibited the highest sequence similarity to it (based on Smith–Waterman alignment scores in MATLAB). These consensus sequences were then used as references to calculate G-to-A mutations within GG and GA dinucleotide motifs in hypermutated sequences.

### SIV sequences

We downloaded all 55,164 available SIV sequences from the LANL HIV databases (accessed 24 Dec 2024). After grouping sequences by subtype and removing duplicates, 39,608 unique sequences remained for analysis. Unlike HIV, hypermutated SIV sequences are not fully annotated in this database. Therefore, we implemented a two-step analysis to accurately identify hypermutated sequences:

*Step 1 (Subtype-wide pairwise screening)* – We used pairwise alignment to align each SIV sequence with all other sequences within the same subtype, regardless of donor origin, using the Smith–Waterman algorithm implemented in MATLAB. For sequence pairs with ≥60% genomic overlap, Hypermut 3^55^ was applied to test for significant G-to-A enrichment in APOBEC-preferred contexts (GpRpD, where R = G or A and D = A, T, or G but not C) at *p* < 0.05. This analysis identified 198 candidate hypermutated sequences, which were subsequently re-evaluated in Step 2 using only sequences from the same donor to confirm true hypermutation status.

*Step 2 (Within-animal confirmation of SIV hypermutation)* **–** To validate the 198 candidate hypermutated SIV sequences, all sequences were grouped by animal, revealing that the candidates originated from 107 animals. For each candidate, 3–5 non-hypermutated sequences from the same animal that shared ≥60% genomic overlap and exhibited the highest sequence similarity to the hypermutated sequence were selected. Candidates with fewer than three overlapping and similar normal sequences were excluded to ensure reliable consensus construction for G-to-A mutation quantification. The selected sequences were multiply aligned using the *multialign* function in MATLAB to generate a within-animal consensus reference. Each candidate hypermutated sequence was then aligned to its corresponding consensus, and hypermutation was reassessed using Hypermut 3.^55^ Of the 198 initial candidates, 171 retained *p* < 0.05 and were confirmed as hypermutated SIV sequences (**Supplementary Table S1**).

### Virus constructs

We used HIV_IIIB-C200_, which is a full-length infectious molecular clone, consisting of the provirus backbone of HXB2 and the SphI-SalI region of pNL4-3 (GenBank EU541617).^56^ Additionally, we obtained SIVmac239 SpX from Dr. Ronald C. Desrosiers (HRP-12249, lot# 150270, BEI Resources).^57^

### A3 expression constructs

We generated a series of A3G expression constructs from an A3G pcDNA3.1(+) base vector with a C-terminal 3xHA tag.^45^ Briefly the human A3G CDS was PCR amplified from its original source^58^ into the Xho/EcoRI sites of pCDNA3.1 with a 5′-HA tag (MYPYDVPDYA) encoded primer. Different introns were inserted into the A3G coding region between exons 1 and 2 to generate the three plasmids: a “Control Intron” plasmid containing the second intron in human hemoglobin subunit gamma 2 (*HBG2*), an “Intron 1” plasmid containing the first intron in rhesus macaque *A3G*, and an “Intron 1 rSNP” plasmid containing the first intron in rhesus macaque *A3G* with a reverted single nucleotide polymorphism (SNP) as discussed in the text. The second intron in *HGB2* was amplified from Jurkat T cell genomic DNA (ATCC TIB-152) using primers 5’-ACC ACC ATG AAG CCT CAC TTC AGG TGA GTC CAG GAG ATG TTT CAG CAC TGT TGC C-3’ and 5’-GAT ACA TTC GCT CCA CTG TGT TTC TGT TGA GAT GAA AGG AGA CAA TAA AGA TGA A-3’ and inserted via Gibson Assembly. The first intron in the rhesus macaque *A3G* was inserted using 4 overlapping gBlocks matching reference sequence NM_001198693.2 via Gibson assembly. The rhesus macaque *A3G* construct containing the reverted ‘predictive’ SNP was generated from prior “Intron 1” construct by site-directed mutagenesis using primers 5’- TCC AGG TCC CCT GCT GAG ACT CTC CCc TGA AAG TCA TGG CTG AGG TGG TGC TCC-3’ and 5’-GGA GCA CCA CCT CAG CCA TGA CTT TCA gGG GAG AGT CTC AGC AGG GGA CCT GGA-3’.

pTwist-CMV-Puro–rhesus A3Gi, is an expression plasmid designed to express canonical rhesus A3G using a codon-optimized construct containing a chimeric intron from the human β-globin and immunoglobulin heavy-chain genes positioned between exons 5 and 6. We removed the sequence between the MulI and PvuII restriction sites of pTwist-CMV-Puro-rhesus A3Gi by digestion and ligated the plasmid backbone to gene blocks designed with the CDS replacing exon1 with the predicted coding sequence of exon1b starting at the first methionine (A3G_M1_) or the second (A3G_M2_). We used whole plasmid sequencing through Plasmidsaurus to confirm the sequence of all viral and expression constructs.

### A3G expression and viral stock production

Chemically competent One Shot TOP10 *E*. *coli* (C404003, Thermo Fisher) was transformed with human and rhesus A3G plasmids (pcDNA3.1(+) backbone) and HIV_IIIB-C200_ plasmid. Chemically competent MAX Efficiency Stbl2 *E. coli* (10268019, Thermo Fisher) was transformed with SIV_mac239_ Spx plasmids (HRP-12249, BEI Reagent Resource, NIAD, NIH). Plasmids were grown on an LB-agarose plate containing 100 μg/ml ampicillin overnight at 37 °C. Colonies were selected and then resuspended in tubes containing 100 μg/ml ampicillin in LB broth for 8 hours. After 8 hours, these cultures were grown at 37°C overnight in a larger flask containing 100 μg/ml ampicillin in LB broth.

For transfection, we plated HEK293T cells (RRID:CVCL_0063) at 2 x 10^5^ cells per well in 2 mL of DMEM Complete (89% DMEM [30-2002, ATCC], 10% FBS [R&D Systems, S11550H], 1% 100x GlutaMax [35050061, Gibco]) in 6-well plates and grew them for 24 hours prior to transfection. Cells were transfected using PolyJet *In Vitro* Transfection Reagent (SL100688, SignaGen Laboratories) with 1 µg of each indicated expression construct. A GFP expression construct was used as a positive control, and transfection efficiency was confirmed using confocal laser microscopy. After 48 hours, cells were washed with PBS/EDTA, released with trypsin (25300054, Gibco), and pelleted by centrifugation. The cell pellets were resuspended in 300 µL/1-2 M cells of HED lysis buffer [25 mM HEPES pH 7.4 [(25060CI, Corning) 15 mM EDTA (15575020, ThermoFisher), 10% glycerol, and Roche cOmplete, mini EDTA-free protease inhibitor cocktail (04693159001, Millipore-Sigma)], then frozen at −80°C until needed. The cells were then lysed in a sonicating water bath at 4°C for 20 min. After sonication the lysate was supplemented by PureLink RNase A (12091021, Thermo Fisher Scientific) to a concentration of 100 μg/mL and placed on a rotating shaker for 1 hour at room temperature. The lysate was then clarified by centrifugation at 18,000 x *g* at 4°C for 10 min.

For growing virus (HIV_IIIB-C200_ and SIV_mac239_ Spx), we plated HEK293T cells at 4 x10^6^ cells/10 mL DMEM Complete in a 100 mm dish 24 hours before transfection. PolyJet was used as the transfection reagent where 10µg of DNA was transfected for 48 hours. After 48 hours, supernatant was collected and syringe filtered (0.45 µM). Virus titer was determined by core antigen for HIV (p24) and SIV (p27) ELISA (5421 and 5436, Advanced Bioscience Laboratories, Inc., respectively).

### Immunoblotting

We ran the whole PBMC lysates and recombinant proteins on a 12% TGX Stain-Free™ FastCast™ SDS-PAGE gel (1610185, Bio-Rad Laboratories) at 70 V for 30 minutes then 120 V for 150 minutes. Proteins were electrotransferred to a Immun-Blot Low Fluorescence PVDF membrane (1620260, Bio-Rad Laboratories) at 90 V for 60 minutes on ice. Membranes were blocked in 5% skim milk in PBST and probed with primary antibody [1:2000 anti-A3G (ARP-10201, BEI Reagent Resource, NIAD, NIH,^59^), 1:10000 anti-HA Tag (3724, Cell Signaling), 1:5000 anti-CYPA (2175, Cell Signaling), 1:1000 anti-RPL13a (2765, Cell Signaling)] at 4°C overnight. Membranes were washed in PBST three times for 10 minutes each. After washing, membranes were incubated with secondary antibody [1:2000 anti-rabbit HRP (7074, Cell Signaling Technologies)] for one hour at room temperature. For stain-free total protein normalization, membrane was imaged using the Stain-Free blot option on a Bio-Rad ChemiDoc MP imager prior to treatment with ECL substrate. For detection of proteins, membranes were incubated with Clarity Max Western ECL Substrate (1705062, Bio-Rad Laboratories) and visualized on a Bio-Rad ChemiDoc MP imager.

The rhesus recombinant A3G protein (pcDNA3-N-terminus-1xFLAG tagged rA3G [NP_001185622]) and the human recombinant A3G protein (pET-N-terminus-6xHis tagged SUMO–hA3G [NP_068594]) were used as controls in the immunoblots.^60^ The human recombinant A3G protein has a replacement fragment (^139^CQKRDGPR^146^ to ^139^AEAG^142^) for solubility enhancement and a PreScission protease cleavage site between SUMO and hA3G. We quantified these recombinant proteins using *RC DC* Protein Assay Kit II (Bio-Rad Laboratories, 5000122) prior to loading. De-identified human PBMCs from two independent, seronegative donors (Hu1 (male) and Hu2 (female)) used in the immunoblots were generously donated by Dr. Schlesinger, and rhesus PBMC samples from five independent, seronegative donors (Rm1 (female), Rm2 (female), Rm3 (female), Rm4 (male), Rm5 (female)) were obtained from the Southwest National Primate Research Center.

### Deaminase assay to determine dinucleotide substrate preference

We incubated HEK293T cell lysates in HED buffer with one of five fluorescently labeled 43-mer substrates (IDT) for 2 hours at 37 °C: 5’-ATT ATT ATT AC (CC/TT/GC/AC/ideoxyU) AAA TGG ATT TAT TTA TTT ATT TAT TTA TTT/36-FAM/-3’. Following a 2-hour incubation period, samples were incubated at 95 °C for 10 minutes to halt enzymatic activity. A UDG mix (10x UDG reaction buffer, UDG [1U/mL], (EN0361, ThermoFisher) and DEPC water) were added to the reaction for a final concentration of 0.1U/reaction UDG) and incubated at 37°C for 10 minutes.

After the incubation with UDG, we added 0.6M NaOH for a final reaction concentration of 100 mM NaOH and incubated at 37°C for 20 minutes. Following incubation with NaOH, reactions were heated to 95 °C for 3 minutes. The solutions were then mixed with 2X RNA loading dye (R0641, ThermoFisher) and run on 15% TBE-urea polyacrylamide gels at 180 V for 90 minutes. The gels were imaged with a Bio-Rad ChemiDoc MP imager.

### Cell line electroporation

We electroporated HEK293T (ATCC CRL-3216) cell lines with either 0.25 or 0.5 µg of each indicated expression construct. For each reaction, 2 x 10^5^ cells were counted and pelleted by centrifugation at 400 x g for 5 minutes. The supernatant was removed by aspiration, and the pellet was resuspended in 20 µL of supplemented room-temperature SE electroporation buffer. The resulting cell suspensions were then mixed with plasmid constructs in TE buffer, pipetted into a 96-well electroporation cuvette, and electroporated using the Lonza 4D-Nucleofector X Unit with pulse code CM-130. Immediately after electroporation, 100 µL of warmed media was added to each cuvette and the cuvette was placed in an incubator at 37°C for 30 minutes to allow cells to recover for 30 minutes. Afterwards, cells were transferred into 12-well culture plates prefilled with 400 µL of warm complete media. The cells were then returned to an incubator at 37°C for two days to allow for protein expression. After two days, cells were resuspended and divided in half to make whole cell lysates and measure viability. Whole cell lysates were prepared by washing cells with 200 µL of 1xPBS and resuspending them in 50 µL of 2.5x Laemmli Sample Buffer prior to protein denaturation by heating at 98°C for 20 minutes. Viability was assessed using the CellTiter-Glo Luminescent Cell Viability Assay. Cells were resuspended in 100 µL of media and transferred to a 96-well white plate. 100 µL of CellTiter-Glo Reagent was added, the plate was covered in foil and set on an orbital shaker for 2 minutes. Following the two minutes, the plate was left to incubate at room temperature for 10 minutes and luminescence was recorded using a FLUOstar Omega Plate Reader.

### CRISPR-Cas9 ribonucleoprotein (crRNP) production

We used a total of 7 CRISPR guide RNA to make crRNPs for knock-out experiments in rhesus macaque PBMCs. Predesigned non-targeting (U-007502-01-20, Dharmacon) and Cyclophilin A (CYPA/PPIA) targeting crRNA (CM-004979-03, Dharmacon) served as controls. Custom crRNA targeting rhesus macaque A3G were designed and ordered from Dharmacon with the following sequences: 5’-CUC UUA UGU CUU CAG AAA CA-3’, 5’-UUA UGU CUU CAG AAA CAU GG-3’, 5’-CAG AAA CAU GGU GGA GCC AA-3’, 5’-ACG AAU GUG CGU GGA UCC AU-3’, 5’-AAG UUG GAG ACG AAU GUG CG-3’. Resulting A3G crRNPs were multiplexed in equal amounts prior to electroporation. To make the crRNPs, lyophilized crRNA and trans-activating CRISPR RNA (tracrRNA) (U-002005-50, Dharmacon) were resuspended at 160 µM in 10 mM Tris-HCL (7.5 pH) with 150 mM KCl. Resuspended cRNA and tracrRNA were combined in equal parts and incubated at 37°C for 30 minutes. The resulting gRNA:tracrRNA complex was combined with 1.6 parts 100 mg/mL poly-L-glutamic acid sodium salt (PGA) (P4761-100MG, Sigma Aldrich) and 2 parts Cas9 (University of California Berkeley MacroLab) in a whisking motion. The resulting solution was incubated at 37°C for 15 minutes, divided into 4 µL aliquots or pooled, and stored at −80°C until use.

### Rhesus macaque PBMC thawing and stimulation

We thawed the rhesus macaque (Rm6, female) PBMCs in a 37°C water bath and added dropwise to pre-warmed RPMI-10 with 50 U/mL Benzonase to prevent cell clumping. RPMI-10 media consisted of RPMI 1640 (Corning #MT15040CM) with 10% Fetal Bovine Serum (Gibco #16140-071), 1% L-glutamine (35050061, Gibco), 1% MEM Nonessential Amino Acid Solution (25025CI, Corning), 2.5% 1M HEPES (SH30237.01, Cytiva), and 1% Penicillin-Streptomycin Solution (SV30010, Cytiva). PBMCs were spun at 300xg for 10 minutes (break low) and rinsed in 25 mL of pre-warmed media. Washed cells were resuspended at 2 x 10^6^ cells/mL in cRPMI with 7µg/mL of Concanavalin A (ConA) (00497893, Invitrogen) and put at 37°C 5% CO2 for 72 hours to recover and stimulate. After 72-hour recovery, cells were spun at 300xg for 10 minutes (break low) and washed again in 25 mL pre-warmed media. Washed cells were resuspended at 2 x 10^6^ cells/mL in RPMI-10 media with 40 U/mL of IL-2 (130097744, Miltenyi Biotec) and returned to a 37°C incubator for 48 hours to further stimulate.

### Rhesus macaque PBMC editing using CRISPR-Cas9

Prior to CRISPR editing, we assessed cell stimulation by anti-CD25 (1130-115-535, Miltenyi Biotec) immunostaining of cell aliquots from unstimulated, ConA stimulated, and ConA + IL-2 stimulated PBMCs. Stimulated PBMCs were counted, centrifuged at 400xg for 5 minutes, and resuspended in P3 electroporation buffer (V4SP-3960, Lonza). A total of 4 electroporation reactions were conducted each consisting of 200,000 cells in 20 µL of electroporation buffer and 4 µL of crRNPs in a 4D-Nucleofector 96-well unit using pulse code EO-115. Electroporated cells were immediately supplemented with 100 µL of pre-warmed RPMI-10 and allowed to recover for 30 minutes in a 37°C incubator. After recovering, edited cells were moved into a 96-well flat-bottom culture plate prefilled with warm media with IL-2 at a final concentration of 40 U/mL. The plate was set to incubate at 37°C 5% CO2 for 72 hours. After 48 hours cells were fed with 50 µL of RPMI-10 with 40 U/mL of IL-2. After the full 72 hours cell pools were split in half to collect whole cell lysates and assess viability by CellTiter-Glo (G7571, Promega).

### Mass spectrometry of whole PBMC lysates

Human PBMCs were either donated or obtained from STEMCELL Technologies (Hu1, male and Hu2, female). Rhesus macaque PBMCs were obtained from the Southwest National Primate Research Center (SNPRC) (Rm1, female and Rm2, female). Aliquots of the PBMCs containing up to 100 µg or protein were lysed in 100 µL 12% phosphoric acid and 10% SDS/50 mM triethylammonium bicarbonate (TEAB). Proteins were then bound to S-Traps (micro; Protifi), reduced/alkylated with a mixture of tris(2-carboxyethyl)phosphine hydrochloride (TCEP) and 2-chloroacetamide, and then digested into peptides for 2 hr at 37°C with trypsin (sequencing grade; Promega) in 50 mM TEAB. Peptides were eluted from the S-Traps sequentially with 50 mM TEAB, 0.2% formic acid, and 0.2% formic acid in 50% aqueous acetonitrile. The pooled eluates were dried by vacuum centrifugation and redissolved in 30 µL of starting mobile phase (see below). Peptides were analyzed by capillary HPLC-electrospray ionization tandem mass spectrometry on a Thermo Scientific Orbitrap Fusion Lumos mass spectrometer. On-line HPLC separation was accomplished with an RSLC NANO HPLC system (Thermo Scientific/Dionex) interfaced with a Nanospray Flex ion source (Thermo Scientific) on a PepSep column [PicoFrit Resolve™ (360 µm OD, 75 µm ID, 10 µm tip, 15 cm, C18 AQ 3.0 µm, 120 Å; New Objective, Inc.)]. Mobile phase A was 0.5% acetic acid (HAc)/0.005% trifluoroacetic acid (TFA) in water and mobile phase B was 90% acetonitrile/0.5% HAc/0.005% TFA/9.5% in water; the gradient ran from 3% to 42% B in 60 min at a flow rate of 800 nL/min.

Precursor ions were acquired in the orbitrap in centroid mode at 120,000 resolution (*m/z* 200); data-dependent higher-energy collisional dissociation (HCD) spectra were acquired at the same time in the linear trap using the “rapid” speed option (30% normalized collision energy). Other MS scan parameters included: mass window for precursor ion selection, 0.7; charge states, 2 – 5; dynamic exclusion, 15 sec (± 10 ppm); intensity to trigger MS2, 50,000. In addition to the standard DDA-MS settings, an “inclusion list” of previously detected human and rhesus A3G peptides was added to the scan strategy with an intensity trigger of 5,000. This enhanced the likelihood of fragmenting peptides of interest if they were present above background. Mascot (v3.1.0; Matrix Science, London UK) was used to search the spectra against a combination of the following databases: a database containing sequences of investigators’ recombinant proteins (799 sequences; 348,419 residues), common contaminants (not including *Bos taurus* proteins, (124 sequences; 62,564 residues) and either UniProt_Macaca_mulatta_ref 9544_20220315 (44,390 sequences; 28,513,169 residues) or UniProt_Human_ref 9606_20220216 (20,588 sequences; 11,394,277 residues), as appropriate for the sample. Peptide tolerance was 20 ppm, fragment tolerance, 0.8 Da. Cysteine carbamidomethylation was set as a fixed modification and methionine oxidation and deamidation of glutamine and asparagine were considered as variable modifications. Trypsin was specified as the proteolytic enzyme, with two missed cleavages allowed. The Mascot search results were imported into Scaffold (version 5.3.4, Proteome Software Inc., Portland, OR). Acceptance of peptide and protein identifications were based on the following: peptides, greater than 95.0% probability by the *Percolator* posterior error probability calculation^61^; proteins, greater than 99.0% probability based on the *Protein Prophet* algorithm.^62^ A minimum of two identified peptides was required. These settings resulted in a protein-level FDR of 0.3%. Proteins that contained shared peptides and could not be differentiated based on tandem-MS analysis alone were grouped to satisfy the principles of parsimony.

### Mass spectrometry of electrophoretic protein bands from PBMCs

We ran whole PBMC lysates from rhesus macaque (Rm1 and Rm2) on two separate 12.5% Criterion™ Tris-HCl protein gels (3450015, Bio-Rad) in tandem. One gel with whole cell lysates was subsequently stained with GelCode Blue Stain (24590, Thermo Fisher Scientific) and then fixed with 10% methanol/7% acetic acid for downstream band excision and mass spectrometry. The other gel was subjected to immunoblotting (as above) to identify the regions for band excision. Recombinant rhesus macaque A3G protein was run on another pair of gels and processed as above as a positive control and to identify which peptides were most abundant mass spectrometry.

We excised the gel bands of interest from the fixed gels and cut into pieces approximately 2 mm x 2 mm followed by addition of 100 µL 12% phosphoric acid and 10% SDS/50 mM TEAB. After pulverization of the gel pieces with a small pestle, the samples were centrifuged at 21000 x *g* for 5 minutes. The supernatants were removed, mixed with additional 10% SDS in 50 mM TEAB, reduced with TCEP, alkylated in the dark with iodoacetamide, and applied to S-Traps micro (ProtiFi) for tryptic digestion (sequencing grade; Promega) for 2 hrs in 50 mM TEAB. Peptides were eluted from the S-Traps sequentially with 50 mM TEAB, 0.2% formic acid, and 0.2% formic acid in 50% aqueous acetonitrile The pooled eluates were dried by vacuum centrifugation and redissolved in 30 µL of starting mobile phase (see below). Equal volumes were analyzed by capillary HPLC-electrospray ionization tandem mass spectrometry on a Thermo Scientific Orbitrap Exploris 480 mass spectrometer. On-line separation was accomplished with a Vanquish

Neo UHPLC system (Thermo Scientific) using a PepSep column (Bruker; ReproSil C18, 15 cm x 150 µm, 1.9 µm beads). Mobile phase A was 0.5% acetic acid (HAc)/0.005% trifluoroacetic acid (TFA) in water and mobile phase B was 90% acetonitrile/0.5% HAc/0.005% TFA/9.5% in water; the gradient was run from 3% to 42% B in 60 min at a flow rate of 0.8 μL/min. Precursor ions were acquired in the orbitrap in centroid mode at 120,000 resolution (m/z 200); EASY-IC™ mode, Run Start; data-dependent higher-energy collisional dissociation (HCD) spectra were subsequently acquired in the orbitrap for up to 20 precursors (normalized collision energy, 27%; resolution 15000). Other MS scan parameters included: m/z window for precursor ion selection, 1.2; charge states, 2 – 5; dynamic exclusion, 20 sec (± 10 ppm); intensity to trigger MS2, 50,000. An inclusion list of A3G peptides was utilized as described above. A3G protein sequences were inferred from whole exome sequencing data for rhesus macaques Rm1 and Rm2, obtained from the Southwest National Primate Research Center (SNPRC). Mascot (v3.1.0; Matrix Science, London UK) was used to search the spectra against a combination of the following databases: a database containing sequences of investigators’ recombinant proteins (799 sequences; 348,419 residues) and SwissProt 2023_01 (569,213 sequences; 205,728,242 residues). Peptide and fragment tolerances were 10 ppm. Other data processing parameters were the same as described above.

### Analysis of a publicly available mass spectrometry dataset from lymphoblastoid cell lines

We reanalyzed a publicly available mass spectrometry dataset of lymphoblastoid cell lines (LCLs) from human, chimpanzee, gorilla, orangutan, and rhesus macaque for A3G-specific peptides.^63^ The spectra were analyzed using Proteome Discoverer (v2.3, Thermo Fisher Scientific) with the Mascot search engine (v2.6, Matrix Science) and searched against a nonredundant database comprised of all UniProt canonical proteomes from five species (human, chimpanzee, gorilla, orangutan, and rhesus macaque).^64^ Peptide relative abundances were normalized as described in the original paper.^63^

### RNA extraction for long-read sequencing

We obtained the olive baboon (*Papio anubis, N* = 3, 2 males) and common marmoset (*Callithrix jacchus, N* = 5, 4 males) PBMCs from the SNPRC. Snap-frozen PBMCs were defrosted and revived in RPMI-10 (RPMI 1640 [MT15040CM, Corning], 10% FBS, 1% L-Glutamine, 1% non-essential amino acids [25025CI, Corning], 25mM HEPES, and 1% Penicillin/Streptomycin [30002CI, Corning) before being processed for RNA extraction using the RNeasy Mini kit (Qiagen). RNA quantity and integrity were measured using the Qubit RNA Broad Range Assay (Q10210, ThermoFisher) and Agilent TapeStation RNA ScreenTape Assay (5067-5576, Agilent Technologies), respectively. Only RNA samples with an RNA Integrity Number (RIN) greater than seven were used for sequencing. Library preparation using the PacBio Kinnex full-length RNA kit and long-read sequencing was performed by Novogene using the PacBio Revio system (Novogene, USA).

### RNA-seq data processing for visualization of A3G splicing

We retrieved RNA-seq datasets from the Sequence Read Archive (SRA) database using sratoolkit (v2.5.7). The quality of datasets was assessed using fastqc (v0.11.8) and multiqc (v1.19). PacBio Iso-seq datasets were quality controlled and processed using standard Iso-seq workflow (https://isoseq.how/). FASTQ files were trimmed and filtered using trimmomatic (v0.39). After quality control, filtered reads were aligned using one of two mappers: STAR (v2.7.10b) and minimap2 (v 2.30) for short- and long-reads, respectively. For STAR, the parameters used were - -outFilterType BySJout --outSAMtype BAM SortedByCoordinate. For minimap2, the parameter - ax splice:hq was utilized. Aligned reads were subsequently filtered using SAMtools (v1.16.1) with the -F 2308 flag to remove unmapped, supplementary and secondary alignments. Reference genomes, RNA-seq SRR IDs and the A3G coordinates used for mapping and visualization are available in **Supplementary File 1**. Sashimi plots were made via ggsashimi (v1.1.5).

### Ribosome profiling data analysis

We conducted a secondary data analysis on a ribosome profiling dataset that consisted of Ribo-seq and matched RNA-seq from LCLs in human (*N* = 4) and rhesus macaque (*N* = 5). These datasets were obtained from the SRA database (PRJNA292112 [rhesus and human], PRJNA214703 [rhesus], and PRJNA122271 [human]). The quality of these datasets was assessed using multiqc (v1.19), and FASTQ files were trimmed and filtered using trimmomatic (v0.39). For Ribo-seq datasets specifically, the first nucleotide on the 5’ end was trimmed since it is an artifact that arises during library preparation.^65^ As an additional quality control of Ribo-seq data, FASTQ files were first aligned to a reference rRNA file^66^ using the BBMap package (v39.01) containing BBDuk which was used to filter rRNA reads that were not completely removed experimentally. Unaligned reads were subsequently used to map Ribo-seq reads to a reference genome using STAR (v2.7.10b). To ensure that the gene transfer format (GTF) file reflected the most relevant rhesus A3G isoforms, we manually incorporated the most abundant isoform (A3G_M2_ – PB.1708.4), identified from Iso-seq long-read sequencing^63^ as corresponding to the A3G_M2_ variant (**Supplementary Table S2**). This step was necessary because A3G_M2_ is not present in the canonical Mmul_10 GTF annotation. A3G_M1_ was also manually added to the annotation.

We used Ribo-TISH (v0.2.7) for data quality control, ensuring most ribosome footprints (ribosome-protected fragments, RPFs) are approximately 30 nucleotides long.^67^ The open reading frame (ORF) prediction was also conducted using Ribo-TISH. To calculate the P-site of each ribosome within the dataset, Ribo-seQC (v1.1) was used. The P-site output was then used for visualization of Ribo-seq counts with matched RNA-seq using ggRibo (v0.3.1).

### Genomic alignment of nonhuman primate A3Gs and single nucleotide polymorphism analysis

We obtained *A3G* gene sequences for various NHPs from the NCBI database within Geneious Prime (v2025.0.3) except for capuchin (*Cebus imitator*) *A3G*, which was obtained from Ensembl. The alignment of *A3G* genes was performed using the ‘Auto’ algorithm of MAFFT (v7.490). A3G protein alignment was performed using the ‘PPP’ standard algorithm of MUSCLE (v5.1). Single nucleotide polymorphism (SNP) analysis was performed using an in-house bioinformatics pipeline. Briefly, to identify SNPs associated with the defective *A3G* splicing phenotype, NHP samples showing abnormal splicing were classified as the defective group. For each genomic position, we compared the nucleotide state in defective samples with that of non-defective individuals. Any site where the defective group consistently carried a different nucleotide than all non-defective samples were considered a candidate SNP associated with the defective *A3G* splicing pattern.

We identified the exonic splicing enhancers (ESEs) using ESEfinder (v3.0).^68^ The query sequences used for ESEfinder consisted of 180 nucleotides corresponding to exon 1b of rhesus macaque (*Macaca mulatta*) and the equivalent region in human A3G. The sequences were as follows: rhesus (5’-TCC AGG TCC CCT GCT GAG ACT CTC CCG TGA AAG TCA TGG CTG AGG TGG TGC TCC CTG CCC TGG GAG GGT GCT CCC TCT GTG TGC TTC CTG CCA TTC CTG AAC TCT GGA ACT GGG AGA ATT GAA CCA AAT TAT GAT TGG AGC AAT GTG GAA TTT TGT TCA CAG AGC TTC GAT GGA TTT TCT-3’) and human (5’-CCC AGG TCC CCT GCT GAG ACT CTC CCC TGA AAG TCA TGG CTG GGG TGG TGC TCC CGG CCC TGG GAG GGT GCT CCC TCT GTG TGC TTC CCG CCA TTC CTG AAC TCT GGA ACT GGG AGA ATT GAA CCA AAG GAT GAG TAG AGC AAG GTG GAA TTT TGT TCA CAG GGC TTT GAT GGA TTT TCT-3’).

Only the highest motif scores exceeding the standard threshold for each motif are reported. Motifs from ESEfinder are as indicated: SRSF1 (Motif 1: (A/G)GAA(G/A)AG-like) and SRSF1 (Motif 2: RGAAGAA (R=A/G)).

### Infection of primary PBMCs for HIV and SIV hypermutation analysis

We stimulated PBMCs from two human donors (Hu3 (male) and Hu4 (female)) and two rhesus macaques (Rm5 (female) and Rm7 (male)) using the same method in the “Rhesus Macaque PBMC Thawing and Stimulation” section. Stimulated PBMCs were infected with 25ng of p24-normalized HIV_IIIB-C200_ or 25ng of p27-normalized SIV_mac239_ Spx per 1x10^6^ cells, respectively (Day 0). However, Hu4 did not have a productive infection as determined by a p24 ELISA and was therefore excluded from subsequent analysis.

On days 4/5 and 6/7 post infection, 700 µL of supernatant was removed, syringe filtered and stored at -80 °C. 280 µL of supernatant was used to extract RNA using QIAmp Viral RNA Mini Kit (52904, Qiagen) with on column DNase treatment but without carrier RNA and eluting in 30 µL of nuclease free water. Additional RPMI-10 + IL-2 medium was added to replace removed supernatant. Cells were harvested, centrifuged at 300 x *g* for 10 min, and lysed at the endpoint (day 6/7) and treated with RNase A solution using the Gentra Puregene cell kit (158722, Qiagen). DNA yield from the final time point was quantified using Nanodrop. Samples were excluded from downstream analysis when there was insufficient RNA or DNA to make a comparison between viral genome and provirus.

### Generation of HIV/SIV PCR amplicon pools for PacBio long-read sequencing

We reverse transcribed RNA using Maxima H Minus Reverse Transcriptase (EP0751, Thermo Scientific) according to the manufacturer’s instructions with minor modifications. Briefly 7 µL of RNA was premixed with 1 µL Oligo(dT) and 1 µL dNTP Mix (SO131 and R0191, Thermo Scientific), incubated at 65°C for 5 min. and then chilled on ice. Subsequently, the total reaction volume was brought up to 20 µL with 4 µL 5xRT buffer, 0.5 µL RNAseOUT (1000840, Invitrogen) 20 U of Maxima H Minus Reverse Transcriptase and nuclease free water. Reactions were incubated at 50°C for 30 min followed by enzyme inactivation at 85 °C for 5 min. To evaluate each RNA sample for potential DNA contamination, another cDNA reaction was carried out without the addition of the reverse transcriptase.

For each virus, we amplified three regions: *gag*, *env*, and near-full-length (NFL). Specifically, for SIVmac239 (reference accession: M33262.1), *gag* (positions:1,143-2,842) and *env* (positions: 6,749-10,230), and for HIV-1 IIIB (reference accession: KJ925006.1), *gag* (positions: 647-2,646) and *env* (positions: 6,574-9,626) were amplified. For NFL, the outer primers for *env* and *gag* were used for each virus. Candidate primer sets were designed using Primer-BLAST and manually curated to avoid guanosine residues at positions corresponding to known A3 target motif, or where necessary, degenerate primers were used. Candidate primers were experimentally evaluated, and final primer sets were selected based on sensitivity, virus specificity, and amplicon length. Primers were synthesized by Integrated DNA Technologies (IDT) as Ultramers containing, from 5′ to 3′, an adapter sequence, a PacBio barcode, and the virus-specific primer sequence (see **Supplementary File 2**). Near full-length PCR was also performed on DNA samples using the gag forward primer and the env reverse primer.

For each sample, three to four 50 µL PCR reaction were carried out with 2 µL of cDNA or 100 ng of extracted DNA with Platinum Superfi II PCR master mix (Invitrogen), and 500 nM forward and reverse primers for each sample (see **Supplementary File 2** for cycling conditions). Minus–RT controls were included for all RNA-derived sample to confirm the absence of DNA contamination, and no-template controls were incorporated in every PCR run to monitor for reagent contamination, while virus constructs served as positive PCR controls.

PCR products were assessed by agarose gel electrophoresis, pooled by sample and purified using the QIAquick PCR purification kit (Qiagen). Purified PCR products were quantified with Qubit dsDNA BR or HS assay (Invitrogen). To maintain separation of nucleic acid source, two distinct amplicon pools were generated: one from vRNA-derived PCR products and one from proviral DNA-derived PCR products. Within each pool, individual PCR products were combined at defined molar ratios such that control samples and endpoint DNA and RNA samples were represented equally.

Amplicon pools were further cleaned and concentrated using SMRTbell cleanup beads (PacBio) at a 1:1 ratio. Libraries were prepared from 500 ng of each cleaned DNA pool following the standard multiplexed amplicon library preparation protocol using the SMRTbell prep kit 3.0 (PacBio). Sample libraries were barcoded, annealed with standard sequencing primer, and bound to SPRQ polymerase using the Revio SPRQ™ polymerase kit (PacBio), then loaded onto Revio SMRT Cells at a 1 RNA-to-DNA samples ratio; sequencing was performed using the Revio SPRQ sequencing kit with a 30-hour movie length.

### A3-mediated hypermutation analysis of PCR amplicon pools from PacBio long-read sequencing

We demultiplexed PacBio raw BAM files using asymmetric barcodes with lima (RRID:SCR_022749) and removed primer sequences in the same step. HiFi BAM files were converted to FASTQ format and aligned to the corresponding HIV or SIV reference genome derived from the plasmid insert using minimap2 with the map-hifi preset (minimap2 -ax map-hifi). Resulting SAM files were converted to BAM, and secondary, supplementary, and unmapped reads were filtered out using samtools with the -F 2308 flag. Filtered BAM files were sorted and indexed with samtools. A3-mediated hypermutation was quantified with Hypermut 3^55^, and the sequence contexts of mutations were determined using an in-house bioinformatics pipeline. Hypermutation signatures obtained from Hypermut 3 are defined by three different contexts: 1-Overall G-to-A (downstream context RD): Counts all G-to-A mutations where the base immediately downstream is R (purine: A or G) followed by D (A, T, or G, excluding C). This captures broad A3 activity while filtering non-preferred contexts. 2- GG-to-AG/AA (downstream context GD): Specifically, GG-to-AG mutations with downstream GD, which captures A3G-specific hypermutation, and 3- GA-to-AA (downstream context AD): GA-to-AA mutations with downstream AD, hallmarking A3D/F/H activity. To assess A3-induced hypermutation differences between HIV and SIV, we quantified three features: a) Hypermutation rate: the fraction of reads classified as hypermutated (by Hypermut 3) out of all mapped reads, b) Hypermutation burden: Ratio of the observed GG-to-AG/AA and GA-to-AA mutations over the total number of potential A3 target sites within a sequence; and c) the relative percentages of the two A3 signatures (GG-to-AG/AA vs. GA-to-AA) within a hypermutated read.

### Analysis of premature stop codons in hypermutated HIV-1 and SIV sequences

We used the reference gene annotations for HIV-1 IIIB and SIVmac239 to identify A3 target motifs (GG and GA) at in-frame coding positions where a G-to-A mutation would generate a premature stop codon. These positions were defined as potential stop-codon sites. For each hypermutated sequence, stop-codon burden was calculated as the number of observed A3-associated stop codons divided by the total number of potential stop-codon sites in that sequence using a custom in-house script.

### Analysis of the mutational profiles in nonhypermutated sequences

For each HIV-1 and SIV sequence that did not meet the Hypermut 3 significance threshold *(p* > 0.05), we calculated the mutation frequencies for all substitutions stratified by dinucleotide context. For each mutation type, the number of observed mutations was normalized by the number of corresponding dinucleotide motifs within the analyzed reference sequence region covered by the query sequence. This yielded context-specific mutation burden that were comparable across sequences of different lengths.

### Statistical analysis

We conducted statistical data analysis using RStudio (v.2026.05.0+218). Prior to running a statistical test, all statistical model assumptions were tested for and met. The continuous outcome variables were compared using a two-sided Wilcoxon rank-sum (Mann-Whitney) test followed by Benjamini-Hochberg correction to obtain adjusted *p*-values. Categorical variables for hypermutation analysis (**Supplementary Table S6**) were assessed for differences using a proportion test (prop.test).

#### Data Availability

Long-read sequencing data reported in this paper will be deposited into the Sequence Read Archive database at the time of publication. All other publicly available RNA-seq SRR IDs used in this study are available in the supplementary materials. All original code has been deposited at GitHub (https://github.com/Diako-Lab/APOBEC3G-Splicing-Defects-in-Nonhuman-Primate-Models-Result-in-Divergent-Viral-Mutational-Profiles) and is publicly available as of the date of publication. Any additional information required to reanalyze the data reported in this paper is available from the lead contact upon request.

## RESULTS

### Distinct A3-associated hypermutation profiles in HIV and SIV genomes

If aberrant splicing of *A3G* in some NHP species was functionally relevant, we would expect reduced A3G-mediated hypermutation patterns in SIV sequences. To compare patterns of A3-induced mutations in HIV and SIV, we analyzed 932,134 HIV-1 and 55,164 SIV sequences from the LANL databases. The HIV-1 dataset includes only clinical isolates and the SIV dataset includes mostly natural isolates such as SIV_sab92018_ and SIV_smmE660_, and as wells as sequences derived from animals experimentally infected with SIV_mac239_ and SIV_mac251_ molecular clones (**Supplementary Figure S1**). These datasets include 6,152 hypermutated HIV-1 sequences and 171 hypermutated SIV sequences, which were identified using Hypermut 3^55^. Hypermutated HIV-1 genomes are distributed among diverse subtypes (**Supplementary Figure S1**). Similarly, hypermutated SIV genomes are detected in both naturally circulating viruses (*n* = 102) and laboratory-derived SIV clones (*n* = 69) (**Supplementary Figure S1**). Most hypermutated SIV isolates derive from animals infected in the wild or in the lab by natural isolates, including species such as the tantalus monkey (*Chlorocebus tantulus*), green monkey (*Chlorocebus sabaeus*), sooty mangabey (*Cercocebus atys*), and patas monkey (*Erythrocebus patas*).

Our analysis revealed distinct G-to-A hypermutation profiles for HIV and SIV. The majority of hypermutated HIV sequences exhibit a dominant GG-to-AG/AA mutation pattern, consistent with A3G-mediated hypermutation (81.4%; *n* = 5,007). In contrast, most hypermutated SIV sequences are predominated by GA-to-AA mutations, characteristic of other A3 enzymes, including A3A, A3B, A3C, A3D, A3F, and stable haplotypes of A3H^45,69–73^ (82.5%; *n* = 141) (Wilcoxon rank-sum tests: *p* < 2.2 x 10^-16^ for GG-to-AG/AA and *p* = 1.1 x 10^-9^ for GA-to-AA) (**Figure 1A**).

**Figure 1.**
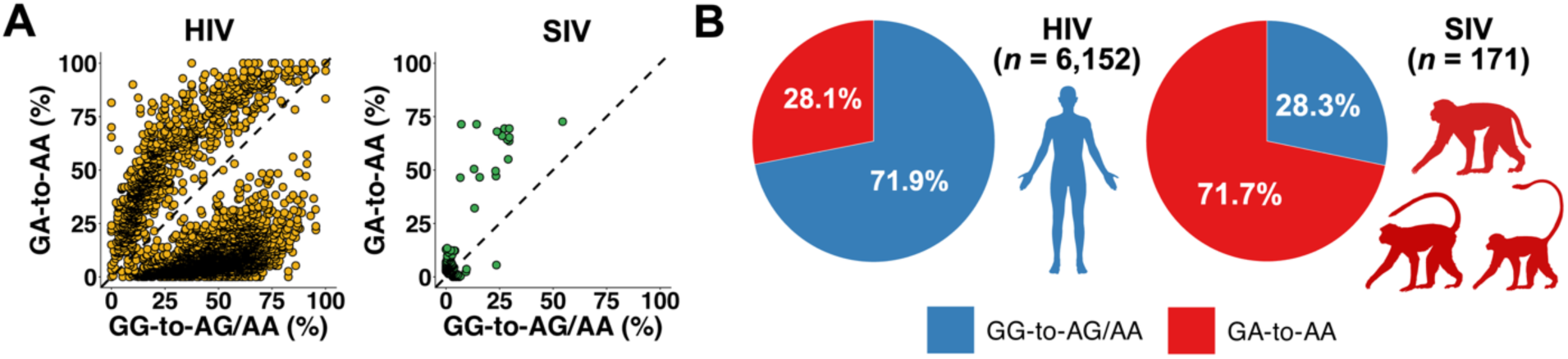
Divergent A3-associated hypermutation profiles in HIV and SIV. **A)** Scatterplots showing the %GG-to-AG/AA mutations (x-axis) versus the %GA-to-AA mutations (y-axis) across hypermutated sequences. Each point represents a viral sequence. Group differences were assessed by a two-sided Wilxocon rank-sum test. **B)** Average mutational signature proportions in HIV and SIV (GA-to-AA vs. GG-to-AG/AA) across sequences.

This disparity is also evident in the overall proportion of GG-to-AG/AA and GA-to-AA mutations across all hypermutated sequences (**Figure 1B**). To ensure that these patterns were not driven by a small number of individuals contributing disproportionate numbers of hypermutated sequences, we performed an additional analysis in which the percentages of each of the two mutational signatures across hypermutated genomes were averaged within each individual, such that each individual contributed only once to the final result. The mutational signature bias remains evident even when sequences are averaged across all donors (**Supplementary Figure S2**). Around 94% (161/171) of NHP hosts carrying hypermutated sequences are macaques (*i.e.*, rhesus, southern pig-tailed, and cynomolgus macaques), with a substantial fraction (85/171) representing experimental SIV_smm_ (sooty mangabey-derived SIV) infections in macaques. Overall, these findings indicate that A3-mediated hypermutation is biased toward GA-to-AA substitutions in NHPs, consistent with reduced A3G-mediated mutagenesis.

It is important to note that, using the HIV/SIV sequences from the LANL database, only the relative proportion of hypermutation signatures can be compared between HIV and SIV, not their absolute burden. This limitation is due to potential biases in sequence deposition, including the historical exclusion of hypermutated sequences because they complicated phylogenetic analyses, the lack of hypermutation-optimized primers for many donor samples, and the comparatively limited annotation and curation of hypermutated SIV sequences. Thus, while LANL sequences provide valuable initial insights into the relative patterns of hypermutation signatures, they do not accurately reflect their true prevalence or magnitude. As described later, to overcome this limitation, we employed a controlled *ex vivo* infection system using primary cells to directly quantify and compare both the relative contribution and overall burden of hypermutation signatures between HIV and SIV.

### Reduced A3G protein levels in rhesus macaque PBMCs

To investigate whether differences in the HIV and SIV hypermutation profiles are due to differences in A3G protein expression levels between humans and rhesus macaques, we performed a western blot on PBMC lysates from four rhesus macaques and two human donors using a cross-species reactive antibody (ARP-10201, BEI Reagent Resource, NIAD, NIH;^59^ **Figure 2A**). As indicated, recombinant rhesus and human A3G proteins used as controls were equally detected by this antibody. In rhesus PBMCs, however, the amount of A3G (normalized to total protein) was substantially lower compared to human PBMCs (see ∼37kDa bands in **Figure 2A**).

**Figure 2.**
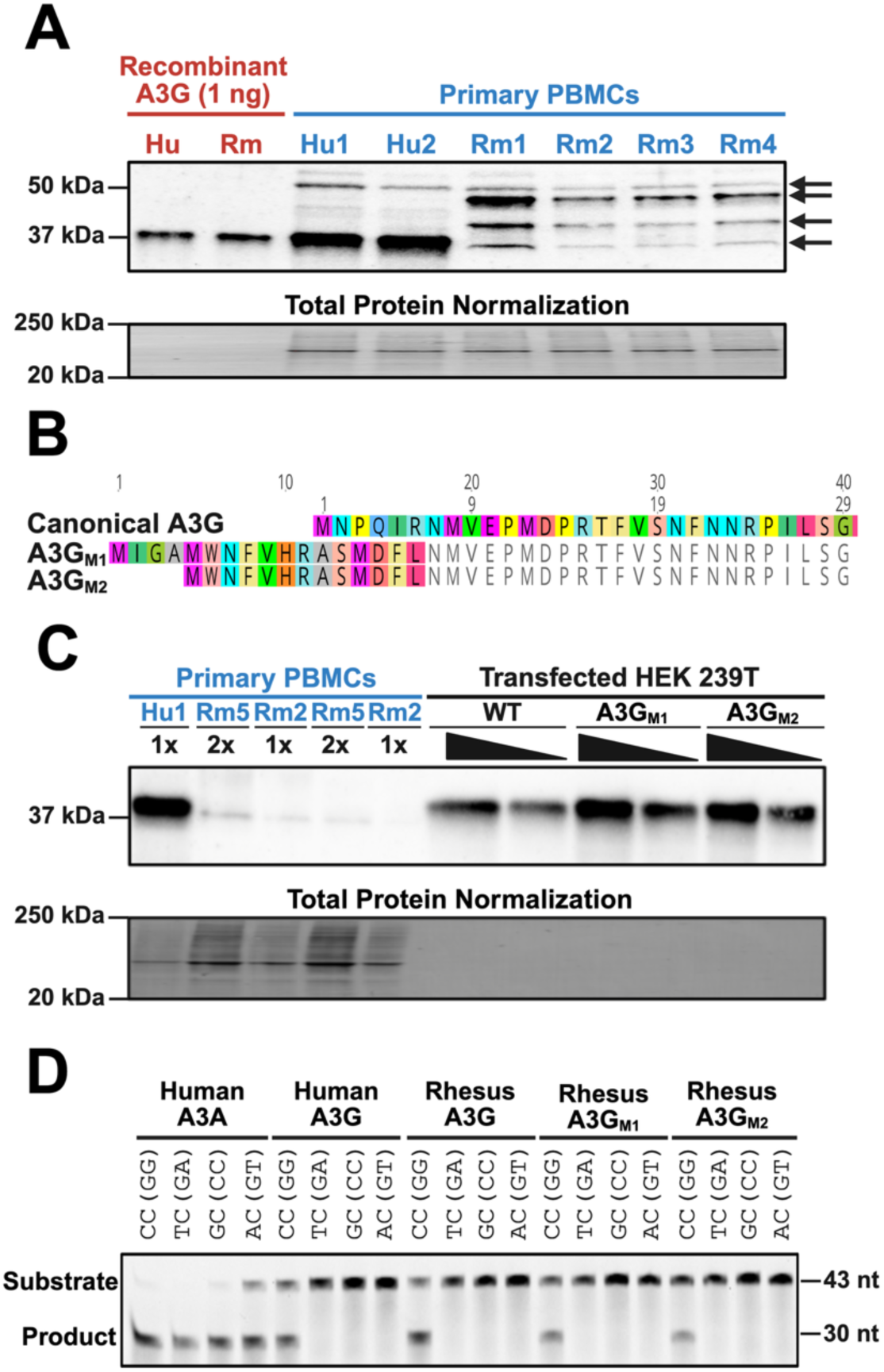
Rhesus macaque PBMCs express less A3G protein than human. **A)** Western blot of human and rhesus recombinant A3Gs and primary PBMCs from two human and four rhesus donors. Hu: Human, Rm: Rhesus macaque. Stain-free total protein normalization was used as a loading control. Arrows indicate bands excised for mass spectrometry analysis. **B)** Partial protein alignment of rhesus canonical A3G, A3G_M1_, and A3G_M2_. **C)** Western blot of primary PBMCs from one human and two rhesus donors alongside transfected rhesus A3G proteins. WT: wild type, canonical A3G. No total protein is shown for transfected cell lysates due to very low amounts of total protein relative to primary PBMC cell lysates. **D)** Deaminase assay gel from HEK239T transfected cell lysates of human A3A (positive control), human A3G, rhesus WT A3G, rhesus A3G_M1_, and rhesus A3G_M2_. Four different oligos containing CC, TC, GC, or AC were used to confirm the preferred motifs.

Notably, rhesus PBMCs displayed two additional bands at ∼40 and ∼50kDa that were not observed in human PBMCs (**Figure 2A**). To determine whether these bands represent novel rhesus A3G proteoforms rather than non-specific antibody binding, we examined rhesus *A3G* mRNA sequences for alternative translation start sites.^74^ If *A3G* translation skips exon 1, initiation could occur at either of two methionines in intron 1, generating two predicted proteoforms: A3G_M1_ (∼46.42kDa; 394 residues) and A3G_M2_ (∼46.05kDa; 390 residues) (**Figure 2B**). A3G_M2_ has previously been predicted from PacBio Iso-seq data^63^ (**Supplementary Figure S3**). We generated expression plasmids for both proteoforms and analyzed them by immunoblotting. Similar to canonical A3G, both A3G_M1_ and A3G_M2_ (∼46kDa) migrated at ∼37kDa, indicating that the additional ∼40kDa and ∼50kDa bands in rhesus PBMCs are not A3G proteoforms.

To confirm that the ∼37kDa band corresponds to rhesus A3G, we knocked out *A3G* in activated rhesus macaque PBMCs using CRISPR-Cas9. Cell activation and viability were comparable across conditions (**Supplementary Figure S4A-B**) and efficient gene editing was confirmed by CYPA knock-out (**Supplementary Figure S4C)**. Immunoblot analysis showed that the ∼40 and ∼50kDa bands were unchanged, whereas the ∼37kDa band was specifically depleted in cells treated with an A3G-targeting guide RNA (**Supplementary Figure S4C**). These results confirm that the ∼37kDa band is rhesus A3G and that the upper bands arise from non-specific antibody binding.

To further validate these findings, we excised these SDS-PAGE gel bands (**Figure 2A**, arrows) and analyzed them by liquid chromatography tandem mass spectrometry (LC-MS/MS) using a data-dependent acquisition (DDA) workflow. No A3G-derived peptides were detected in any of the bands. As a positive control, LC-MS/MS analysis of recombinant rhesus A3G identified 15 unique peptides (76 total) demonstrating that A3G can be readily detection when present at sufficient abundance. We therefore performed DDA LC-MS/MS on whole cell lysates from rhesus and human PBMCs. Two unique A3G peptides and seven unique A3C peptides were detected in the human PBMCs, whereas no A3G peptides were detected in the rhesus PBMCs (**Supplementary Files 3 and 4**).

Although DDA LC-MS/MS is well suited for protein identification, its stochastic sampling limits quantitative comparison across samples. We therefore analyzed a publicly available isobaric-labeling-based LC-MS/MS dataset of lymphoblastoid cell lines (LCLs) from humans, chimpanzees, gorillas, orangutans, and rhesus macaques.^63^ This analysis identified two rhesus A3G peptides, one of which (“IYDDQGR”) was quantifiable across all five species. Among the five species, this peptide was least abundant in rhesus macaques with approximately four-fold lower compared to human (**Supplementary Table S3**). Together, these MS data support our immunoblotting results and demonstrate that A3G protein level is significantly lower in rhesus macaques compared to humans and other primates.

Reduced levels of A3G in rhesus macaque PBMCs would be consistent with the lower frequency of GG-to-AG/AA mutations observed in SIV genomes. An alternative explanation, however, is that rhesus A3G, if induced under certain conditions, might exhibit a different dinucleotide preference (for example, favoring GA-to-AA mutations instead of GG-to-AG/AA). To test this possibility, we performed *in vitro* deaminase assays using single-stranded DNA substrates containing cytidine in different dinucleotide contexts. Lysates from HEK293T cells expressing human A3A (control), human A3G, rhesus A3G, as well as predicted rhesus A3G_M1_, and A3G_M2_ were incubated with each substrate, followed by uracil DNA deglycosylase (UDG) and alkaline treatment to cleave at sites of deamination. Products were resolved by gel electrophoresis. Human A3A displayed broad activity and induced mutations across all tested sequence contexts, whereas both human A3G and rhesus A3G showed a strong preference for GG-to-AG editing. The predicted rhesus proteoforms A3G_M1_ and A3G_M2_ likewise preferentially induced GG-to-AG mutations (**Figure 2D**). Thus, if expressed, rhesus A3G and the predicted proteoforms would have the capacity to induce GG-to-AG/AA mutations in the SIV genome. However, our western blots and mass spectrometry data indicate that rhesus A3G is expressed at very low levels, and there is no evidence for the expression of the predicted A3G proteoforms in rhesus PBMCs.

### Defective splicing of A3G mRNA in nonhuman primates

To elucidate whether differences in A3G protein expression are due to differential splicing of *A3G* between human and NHPs, we analyzed several short-read and long-read RNAseq datasets from diverse NHPs (**Supplementary File 2**) and produced Sashimi plots to identify species-specific splicing patterns in *A3G*. Higher primates are classified into Platyrrhini (New World monkeys, NWMs), which include marmosets, tamarins, capuchins, and squirrel monkeys, and Catarrhini (Old World monkeys, OWMs), which comprise the great apes (humans, chimpanzees, gorillas, orangutans, and bonobos). OWMs are further divided into two subfamilies: Cercopithecinae, including rhesus macaques, baboons, and African green monkeys, and Colobinae, which includes snub-nosed monkeys, colobus monkeys, and langurs (**Figure 3**). In humans and other great apes, *A3G* mRNA has eight exons. However, the Sashimi plots indicated nine exons in *A3G* mRNA from all species in the Cercopithecinae subfamily of OWMs. This additional exon (referred to as exon 1b hereafter), is an intronic region between exons 1 and 2 (**Figure 3**). Interestingly, neither the Colobinae subfamily of OWMs nor NWMs display this alternative splicing event. Analysis of RNA-seq datasets from ten additional rhesus macaques across independent studies further confirmed that inclusion of exon 1b in *A3G* mRNA is highly common and occurs across all tested tissues, including whole blood, CD8+/CD4+ T cells, kidney, cerebellum, and spleen (**Supplementary Figure S5**). Examination of exon 1b revealed that the two PTCs within this region are present in all other Cercopithecinae subfamily members (**Supplementary Figure S6**), indicating that inclusion of exon 1b reduces the expression of a functional A3G across all of these species.

**Figure 3.**
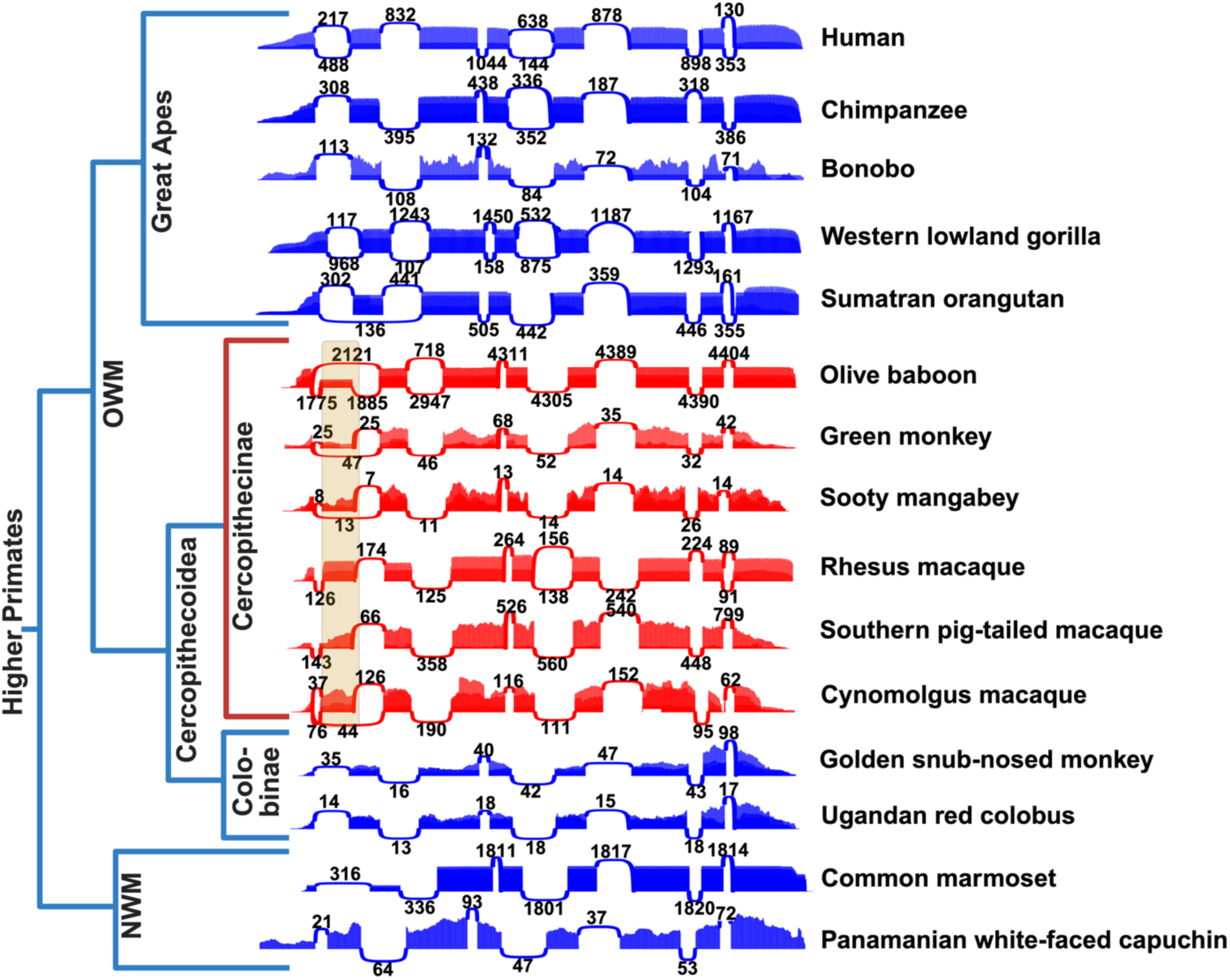
*A3G* mRNA splicing is defective in the Cercopithecinae. Sashimi plots depicting *A3G* splicing across human and NHPs in both Old World monkey (OWM) and New World monkey (NWM) species. Human (*Homo sapiens*; *N = 5*). Chimpanzee (*Pan troglodytes*; *N* = 5). Bonobo (*Pan paniscus*; *N = 2*). Western lowland gorilla (*Gorilla gorilla gorilla*; *N = 3*). Sumatran orangutan (*Pongo abelii*; *N = 5*). Olive baboon (*Papio anubis*; *N = 5*). Sooty mangabey (*Cercocebus atys*; *N = 5*). Green monkey (*Chlorocebus sabaeus*; *N = 5*). Rhesus macaque (*Macaca mulatta*; *N = 5*). Cynomolgus macaque (*Macaca fascicularis*; *N = 5*). Southern pig-tailed macaque (*Macaca nemestrina*; *N = 5*). Golden snub-nosed monkey (*Rhinopithecus roxellana*; *N = 5*). Ugandan red colobus (*Piliocolobus tephrosceles*; *N = 5*). Common marmoset (*Callithrix jacchus*; *N = 5*). Panamanian white-faced capuchin (*Cebus imitator*; *N = 1*). Median values for splice junctions are reported. Yellow shaded area indicates A3G exon 1b presence.

### Ribosome stalling in exon 1b of rhesus A3G

To further investigate the hypothesis that inclusion of exon 1b in rhesus macaque *A3G* mRNA leads to a translational block, we analyzed a publicly available ribosome profiling (Ribo-seq) dataset^75^ alongside a matched RNA-seq dataset^76,77^ from human and rhesus macaque lymphoblastoid cell lines (LCLs). As expected, we found no evidence for retention of the exon 1b region in human *A3G* transcripts in the RNA-seq dataset nor for translation of the exon 1b region of human A3G in the Ribo-seq dataset (**Figure 4A**). However, both the RNA-seq and Ribo-seq datasets from rhesus macaque cells showed reads mapping to rhesus exon 1b, supporting exon 1b inclusion in *A3G* mRNA and its subsequent translation (**Figure 4B**).

**Figure 4.**
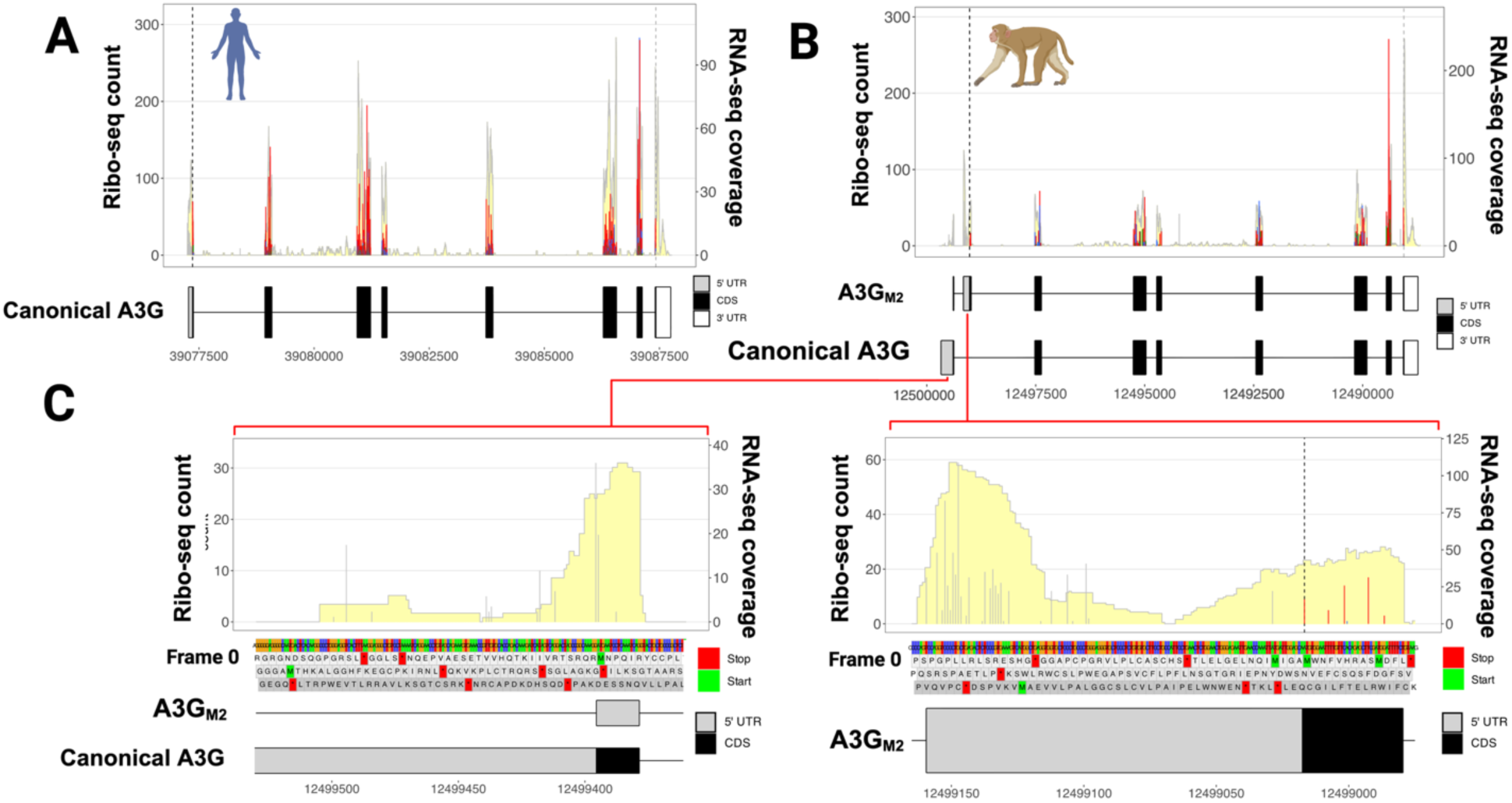
Rhesus *A3G* Exon 1b displays active translation and ribosome stalling. **A)** Pooled Ribo-seq counts and matched RNA-seq coverage of human *A3G* displaying normal and active translation (*N* = 4). **B)** Pooled Ribo-seq counts and matched RNA-seq coverage of rhesus macaque *A3G* displaying lower overall expression and translation (*N* = 5). (C) Pooled Ribo-seq counts and matched RNA-seq coverage of rhesus macaque *A3G* exon 1 (left) and exon 1b (right) displaying lower overall expression and translation (*N* = 4). RNA-seq coverage is shaded in yellow and Ribo-seq counts are determined by P-site calculation and are displayed as grey (not in the main annotated coding sequence (CDS)), red (frame 0), blue (frame 1), or green (frame 2) peaks.

Analysis of the Ribo-seq reads at the start of canonical exon 1 reveals reads supporting translational initiation at the predicted start codon with an expected accumulation of ribosomes at the translation start site (**Figure 4C**, left). However, there is also a marked accumulation of Ribo-seq reads at the start of exon 1b, indicative of ribosome stalling in that region likely due to the presence of the two PTCs (**Figure 4C**, right). This stalling is also reflected in markedly lower Ribo-seq RPKM (reads per kilobase of transcript per million) and TPM (transcripts per million) values for rhesus macaque *A3G* when compared to human *A3G* (**Supplementary Table S3**). Additionally, as expected, other exons in human *A3G* exhibit codon periodicity (i.e., strong enrichment of Ribo-seq reads every third nucleotide) (**Supplementary Figure S7A-C**) while rhesus *A3G* exon 2 does not (**Supplementary Figure S7D**).

It is worth noting that a limited number of ribosome reads are detected downstream of the stop codons at the end of exon 1b, suggesting the presence of a potential secondary translational start site that could give rise to the A3G_M1_ or A3G_M2_ proteoform. To further assess translational initiation sites and predicted translated ORFs based on the Ribo-seq data, we used the Ribo-TISH toolkit, which predicted the translation of A3G_M1_, but not the canonical A3G or A3G_M2_ ORFs (**Supplementary Table S4**).

Overall, our Ribo-seq analyses indicate that the predominant rhesus macaque *A3G* mRNA transcript incorporates exon 1b, creating a small upstream ORF (uORF) spanning exons 1 and 1b ending at a PTC in exon 1b. Downstream of this, a second translational start site could potentially generate a novel proteoform (A3G_M2_), but our immunoblots combined with mass spectrometry analyses show little to no evidence for the expression of A3G_M2_ or any other proteoforms, including the canonical A3G, in rhesus macaque PBMCs.

### Genetic basis of defective A3G mRNA splicing in Cercopethicinae subfamily

Alternative splicing commonly arises from either simple nucleotide polymorphisms or insertions of endogenous elements, such as *Alu* elements, or a combination of both.^78–80^ These primate-specific retrotransposons frequently insert into intronic regions and promote alternative splicing.^81^ They can create pseudo-exons or alter splice site recognition, with antisense *Alu* elements particularly effective due to their poly(A) tails forming robust polypyrimidine tracts adjacent to acceptor sites, enhancing exon inclusion.^82^ Such mechanisms have expanded proteome diversity in primates and humans.^83^ To determine whether these elements or other sequence variants underlie defective *A3G* splicing in Cercopithecinae, we performed a cross-species analysis of the *A3G* locus. (**Figure 5A**). We first examined the potential role of Alu insertions. Consistent with previous findings in northern pig-tailed macaque (*Macaca leonina*),^27^ we identified an *Alu* element (*Alu*YRa1) in the first intron of *A3G* in rhesus macaque, southern pig-tailed macaque, and cynomolgus macaque, but not in other Cercopithecinae species that exhibit A3G splicing defects, including sooty mangabey, green monkey, and olive baboon (**Figure 5A**). Furthermore, NWMs harbor a distinct Alu insertion (*Alu*Sq2) at the same genomic region but maintain normal *A3G* splicing. Collectively, these results indicate that the presence of an *Alu* element alone is insufficient to explain aberrant *A3G* mRNA splicing across Cercopithecinae.

**Figure 5.**
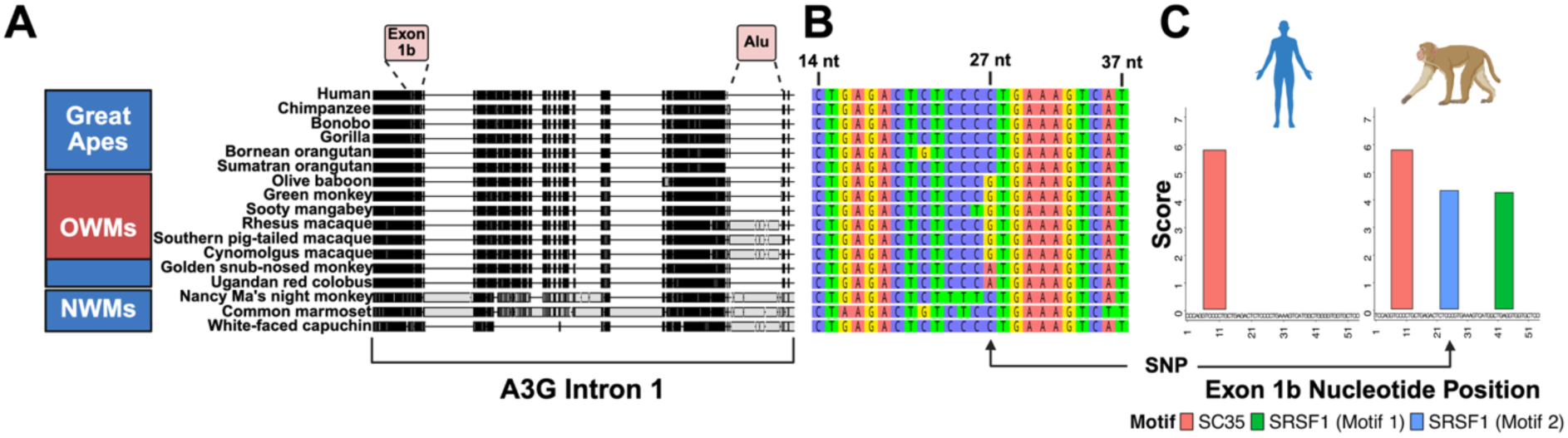
The ‘predictive’ SNP within exon 1b confers a novel exonic splicing enhancer site in rhesus macaque. **A)** *A3G* alignment showing that some Old World monkeys display aberrant *A3G* splicing but do not have the Alu element, and New World monkeys displaying a normal *A3G* splicing, despite having an Alu element in the same genomic location. **B)** Exon1b alignment showing a SNP at position 27 (with respect to the start of exon1b) that is a perfect predictor of *A3G* splicing defect. **C)** Exonic splicing enhancer (ESE) motif scores in human exon and rhesus macaque 1b regions.

We next conducted a cross-species association analysis to investigate whether genetic polymorphism within the *A3G* locus could predict mRNA splicing differences among NHPs. This analysis identified multiple associated SNPs, including one located 27 nucleotides into exon 1b, which is a perfect predictor of aberrant *A3G* mRNA splicing. Presence of guanine (G) at this position correlates with defective *A3G* splicing, whereas the presence of either cytosine (C) or adenine (A) at this position correlates with normal splicing (**Figure 5B**). Notably, this SNP creates a motif recognized by the essential splicing factor SRSF1 (**Figure 5C**). Analysis of exonic splicing enhancer (ESE) sites in rhesus macaque exon 1b revealed a high SRSF1 (Motif 1) and SRSF1 (Motif 2) score at the start of exon 1b, precisely where the SNP resides. These are notably absent in the human ortholog of exon 1b (**Figure 5C**). Together, these data suggest that this SNP may contribute to defective *A3G* splicing in the Cercopithecinae lineage by generating or strengthening a splicing-enhancer motif.

To further investigate the role of this SNP, we generated a series of human A3G expression plasmids containing the mature human *A3G* mRNA reference sequence with rhesus macaque intron 1 inserted into its orthologous site. The SNP was then reverted from a ‘G’ to a ‘C’ (reverted SNP; rSNP). As controls, we also generated a construct lacking an intron and a construct containing a previously reported intron that supports efficient splicing from human hemoglobin subunit gamma 2 (*HBG2*) (**Figure 6A**). Nucleofection of constructs into HEK293T cells was performed with cell viability monitored by a luminescence-based ATP assay and protein expression assessed by western blotting at 48 hours post transfer. Cell viability remained comparable across all conditions (**Figure 6B**). The control A3G expression plasmid lacking an intron expressed high levels of A3G in a dose-dependent manner (**Figure 6C**). Inclusion of the control intron reduced A3G expression slightly. On the other hand, intron 1 from rhesus macaque resulted in near complete suppression of A3G protein expression regardless of dose. Reversion of the SNP slightly increased A3G expression in a dose-dependent manner, though not to the same extent as the control intron. In addition to these experiments, we rearranged (scrambled) the *Alu* element sequence to remove any potential cryptic splice sites and determine whether this can rescue A3G protein expression, however no rescue was observed (**Supplementary Figure S8**). These findings are consistent with the repressive effect of rhesus macaque intron 1, though they suggest additional factors beyond the associated SNP may contribute to aberrant protein expression, at least in this reductionist model.

**Figure 6.**
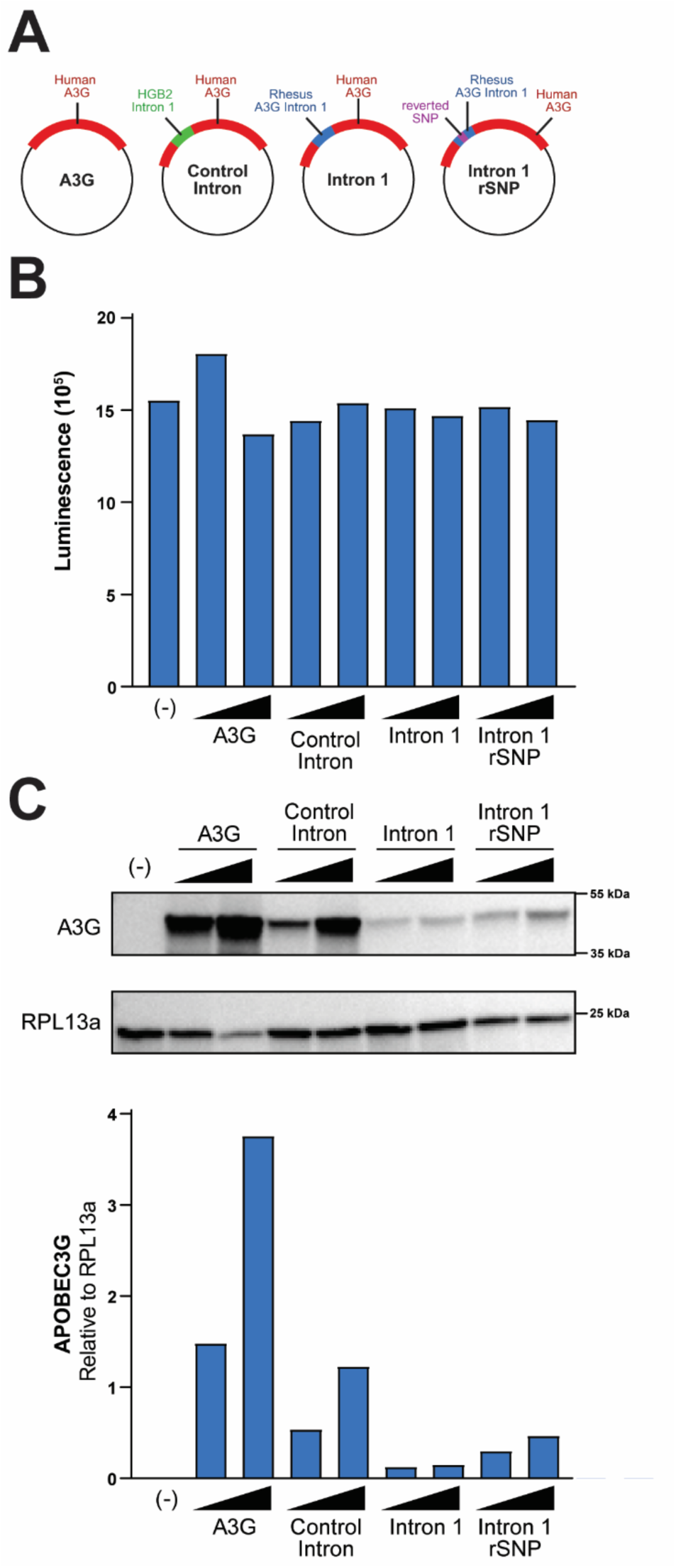
Reversion of the ‘predictive’ SNP within rhesus exon1b partially rescues A3G expression. **A)** Schematic of plasmid constructs, **B)** Cell viability of HEK293T cells expressing constructs assessed by CellTiter-Glo. **C)** Immunoblots of HEK293T cells expressing indicated plasmid constructs, and quantification relative to loading control RPL13a, showing that rhesus intron 1 strongly ablates A3G expression, while reversion of the ‘predictive’ SNP with this intron partially rescues A3G expression.

### Functional impact of differential HIV and SIV mutation profiles driven by A3G splicing differences between human and NHP models

Our hypermutation profiling of HIV and SIV sequences in the LANL databases showed that HIV hypermutation is predominantly characterized by GG-to-AG/AA substitutions, which frequently generate stop codons. In contrast, SIV hypermutation is primarily driven by GA-to-AA substitutions, which are much less likely to create stop codons (**Figure 1**). As a result, we hypothesized that A3-induced mutations are substantially less lethal and are more likely to contribute to the evolution of SIV compared to HIV. To test this hypothesis, we performed two analyses. First, we assessed the molecular source of hypermutated sequences in the LANL databases. Strikingly, while all ∼6,000 hypermutated HIV sequences are from proviral DNA, hypermutated SIV sequences are from both proviral DNA and genomic RNA (**Supplementary Table S1**), suggesting that some hypermutated SIV genomes remain replication-competent and can produce virions.

Second, to recapitulate these results *ex vivo*, we performed a 6/7-day spreading infection experiment using primary human *(N =* 1) and rhesus macaque *(N* = 2) PBMCs infected with HIV-1_IIIB_ and SIV_MAC239_ molecular clones, respectively. Cellular DNA and supernatant RNA were collected at several timepoints post-challenge and near-full-length viral genomes as well as regions within *gag* and *env* were amplified using primers specifically designed to avoid A3 target motifs. Amplicons were sequenced on the PacBio platform, and hypermutated reads were identified and quantified using Hypermut 3 and an in-house bioinformatics pipeline. This analysis generated 1,387,250 HIV (*N* = 1 donor) and 3,897,637 SIV (*N* = 2 animals) reads, including 18,236 and 44,796 hypermutated sequences for HIV and SIV, respectively. To our knowledge, this is the first analysis of hypermutation at such high depth, enabling accurate profiling of mutational signatures.

Analysis of all mutation types across unique reads showed that 65.3% of HIV DNA mutations were G-to-A substitutions, of which 91.6% occurred within the A3 target motifs GG and GA. Similarly, in SIV DNA, 59.7% of mutations were G-to-A changes, with 87.5% occurring in GG and GA contexts. For HIV RNA and SIV RNA, approximately 30% (23.5% HIV; 32.7% SIV) of all single-base substitutions were G-to-A changes, and among these, 75.9% and 77.5%, respectively, occurred within A3-preferred motifs (**Figure 7A**). Collectively, these data indicate that A3-induced cytidine deamination represent the major source of mutations in both proviral DNA and viral RNA.

**Figure 7.**
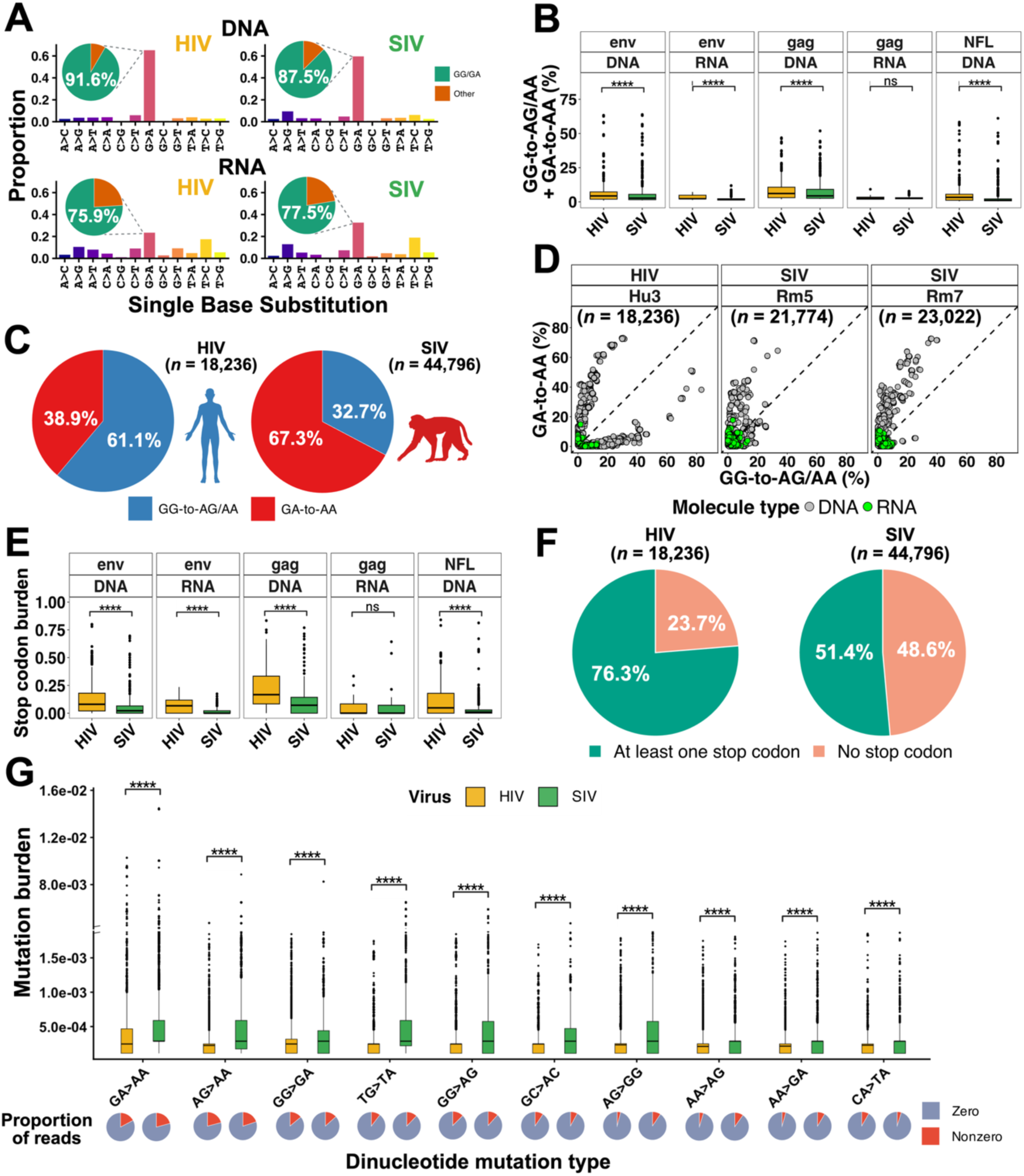
*Ex vivo* infection of rhesus and human PBMCs confirms that A3G-induced hypermutation signature is dominant in HIV, but not in SIV. **A)** Proportion of single base substitutions by virus and molecule type. HIV DNA (*n* = 1,607,749), SIV DNA (*n* = 2,620,240), HIV RNA (*n* = 444,205), SIV RNA (*n* = 1,743,175). Pie charts are dinucleotide substitutions (GG or GA vs other). **B)** A3 mutational burden (GG-to-AG/AA and GA-to-AA) represented by box and whisker plots segregated by virus, molecule type, and PCR region. Lines in box and whisker plots represent the median value. **C)** Average mutational signature proportions in HIV and SIV (GA-to-AA vs GG-to-AG/AA) by sequence. **D)** Scatterplots by donor showing the %GG-to-AG/AA mutations (x-axis) versus the %GA-to-AA mutations (y-axis) across hypermutated sequences, colored by molecule type. Each point represents a single hypermutated virus (panels B and D). **E)** A3-induced stop codon burden represented by a box and whisker plot segregated by virus, molecule type, and PCR region. **F)** Proportion of hypermutated HIV and SIV sequences containing at least one stop codon or no stop codon. Proportion differences in the sequences with ≥1 A3-associated stop codon were assessed using a two-sided two-sample proportion test. **G)** Box and whisker plot of the mutation burden within the top ten dinucleotide contexts in nonhypermutated HIV (*n* = 450,095) and SIV (*n* = 1,548,975) sequences. Pie charts represent the proportion of reads with zero mutations. Group differences were assessed by a two-sided Wilxocon rank-sum test and was corrected by Benjamini-Hochberg procedure (**** = *p*.adj < 0.0001; *** = *p*.adj < 0.001; ** = *p*.adj < 0.01; * = *p*.adj < 0.05; ns = *p*.adj ≥ 0.05).

To accurately assess differences in the A3-driven mutagenesis between HIV and SIV, we considered five complementary features: a) hypermutation rate, defined as the proportion of hypermutated reads among all mapped reads; b) hypermutation burden, defined as the mean fraction of mutated sites per read normalized to the total number of A3 target motifs; c) hypermutation context, defined as the proportions of GG-to-AG/AA and GA-to-AA mutations per hypermutated read, each normalized to the number of GGD and GAD (D: A, G, T) sites, respectively, d) stop codon burden, defined as the number of observed A3-associated stop codons in each hypermutated sequence divided by the total number of potential stop-codon sites; and e) mutational profile of nonhypermutated sequences, defined as the spectrum of normalized mutation frequencies across all mutation types within a dinucleotide context.

The hypermutation rates were 1.46% (17,727 of 1,212,034) for HIV DNA and 2.10% (37,837 of 1,797,608) for SIV DNA. The absolute difference (0.64%), although minimal, was statistically significant (χ²(1) = 1648, *p* < 2.2^-16^). The rate of RNA hypermutation was significantly lower than the rate of DNA hypermutation for both HIV (0.29%) and HIV RNA (0.33%) (**Supplementary Table S6**). This is consistent with frequent viral inactivation following A3-mediated editing. For both DNA and RNA reads (except for *gag* RNA, which showed no difference), the A3-induced mutation burden of HIV was higher than that of SIV (**Figure 7B)**. Additionally, consistent with the analysis of LANL HIV and SIV sequences (**Figure 1A**), the *ex vivo* results demonstrated a clear sequence context distinction between HIV and SIV mutations. In HIV, both GG-to-AG/AA and GA-to-AA hypermutation signatures were observed, with GG-to-AG/AA being more prevalent, whereas SIV sequences predominantly exhibited the GA-to-AA hypermutation signature (**Figure 7C & 7D**).

Hypermutated sequences from both HIV and SIV exhibited a broad range of stop-codon burdens, from zero (no stop codons) to 0.84, indicating that up to 84% of potential GG/GA stop-codon sites were converted to stop codons. Except for *gag* RNA, which showed no significant difference, stop-codon burden was consistently lower in SIV than in HIV across all datasets analyzed (**Figure 7E**). This difference was also evident when examining the proportion of hypermutated sequences lacking A3-induced stop codons. Approximately 49% of hypermutated SIV sequences contained no stop codons, compared with only 24% of hypermutated HIV sequences (two-sided two-sample proportion test, difference = −24.9, 95% CI [-25.7,-24.1], *p* < 2.2 x 10^-16^) (**Figure 7F**). These data strongly indicate that SIV is substantially less likely than HIV to acquire lethal nonsense mutations.

As expected, hypermutated sequences dominated by a GG-to-AG/AA signature were much more likely to be defective, with approximately 93.5% of sequences carrying at least one stop codon. In contrast, only approximately 39.3% of sequences dominated by a GA-to-AA signature contained at least one stop codon (**Supplementary Fig. S9**).

The analyses described above focused exclusively on hypermutated sequences, which contain multiple A3-induced mutations and therefore display a readily detectable A3 signature. In contrast, most viral sequences are classified as nonhypermutated because they harbor relatively few mutations, making A3 activity difficult to infer at the level of individual sequences. We reasoned, however, that if A3 mutagenesis contributes broadly to viral evolution, its signature should become apparent when mutations are analyzed collectively across large numbers of nonhypermutated sequences.

To test this hypothesis, we analyzed only nonhypermutated HIV and SIV sequences and quantified the burden of all mutation types within dinucleotide contexts. Remarkably, GA-to-AA mutations, the hallmark of non-A3G activity, exhibited the highest mutation burden among all mutation classes and were significantly more frequent in SIV than in HIV (**Figure 7G**).

Together, these findings indicate that A3-mediated mutagenesis is a major source of genetic variation not only in hypermutated genomes but across the broader HIV and SIV populations. They further demonstrate that the A3G splicing defect in rhesus macaques and related NHP models functionally reshapes SIV mutagenesis, reducing the generation of lethal stop codons while promoting viral diversification relative to HIV.

## DISCUSSION

Here, we used a multiomics approach to determine the scope, molecular basis, and functional consequences of *A3G* splicing variation across primate lineages. Our study reveals a lineage-specific splicing defect in *A3G* in the Cercopithecinae subfamily of OWMs, which includes widely used NHP models such as rhesus macaques, baboons, sooty mangabeys, and African green monkeys. This splicing defect leads to markedly reduced A3G expression and reshapes the mutational profile of SIV in these species. Given that the preferred mutational context of A3G often results in the generation of PTCs and defective proviruses, reduction of this mutational pressure results in a lower burden of deleterious mutations. This fundamental difference in innate antiviral immunity between commonly used NHP models and humans alters the mutational pressures imposed on replicating viruses, with implications for drug resistance and immune evasion.

The *A3G* defect arises from aberrant splicing that incorporates an intronic region resulting in two PTCs in the *A3G* transcript that cause ribosome stalling, significantly reducing the expression of full length A3G protein. Although not confirmed by mass spectrometry, Ribo-seq data indicate limited amounts of reinitiation occurring at two downstream sites (A3G_M1_ and A3G_M2_) due to a short upstream open reading frame (uORF). Although short uORFs (<40 codons) can permit partial reinitiation, they do not fully protect transcripts from nonsense-mediated decay (NMD),^84–87^ consistent with the persistently low levels of full-length A3G protein observed in primary rhesus macaque cells.

Consistent with Zhang *et al.*’s^27^ findings in northern pig-tailed macaques, we identify *Alu*YRa1 insertion in *A3G* across all macaque species studied. However, our cross-species genomic analysis shows that the presence of an *Alu* alone does not predict splicing defects across Cercopithecinae. Some OWMs lacking *Alu*YRa1, including baboons and African green monkeys, still exhibit aberrant *A3G* splicing, while NWMs retain normal splicing despite harboring an *Alu*Sq2 at the same locus. Instead, we identify a single SNP within exon 1b that correlates with defective splicing and generates a high-scoring SRSF1-responsive exonic splicing enhancer site absent in humans. Our functional assays using chimeric constructs show that rhesus intron 1 represses A3G protein expression, and SNP reversion partially restores it, indicating this variant contributes to, but is not solely responsible for the defect. Together these findings support a model in which lineage-specific *cis*-acting elements drive exon 1b inclusion. Intronic Alu elements may contribute to splicing regulation, but they are not the primary drivers of the defect. The incomplete rescue upon SNP reversion further suggests cooperative interactions among additional *cis*- and *trans*-acting factors likely underlie the *A3G* splicing defect in Cercopithecinae.

Evolutionary analyses suggest that the *A3G* splicing defect arose approximately 15–20 million years ago in the common ancestor of Cercopithecinae and is absent in Colobinae, great apes, and NWMs.^88,89^ The long-term persistence of this defect implies that reduced A3G expression is neither strongly deleterious nor incompatible with effective antiviral immunity in this lineage. This persistence suggests either relaxed constraint on A3G function in this lineage or a lineage-specific evolutionary trade-off favoring reduced mutagenic activity. Notably, signatures of positive selection driven by SIV Vif are evident in A3G across multiple Cercopithecinae species,^90–92^ suggesting that residual A3G activity may still exert selective pressure on viral antagonists. Certain cell types or physiological conditions *in vivo* may promote exclusion of exon 1b. This could generate localized A3G expression that is not apparent in PBMCs. Reduced or targeted A3G activity may represent a lineage-specific evolutionary trade-off that limits the mutagenic costs associated with A3G-mediated genome editing.

Consistent with this prediction, our viral hypermutation analyses reveal clear functional consequences of A3G deficiency in NHPs. Analysis of more than 980,000 HIV-1 and SIV sequences from the LANL database revealed distinct hypermutation patterns. HIV-1 hypermutation is dominated by A3G-mediated GG-to-AG/AA changes that frequently generate deleterious stop codons. In contrast, SIV mutations are predominantly GA-to-AA, a signature associated with other A3 enzymes that is much less likely to create stop codons and is therefore less deleterious.

Analysis of more than 5 million HIV and SIV sequences including over 63,000 hypermutated genomes generated, for the first time, in infectivity experiments using primary human and rhesus PBMCs, not only recapitulated these findings, but also revealed A3 mutagenesis as a major source of HIV/SIV genetic variation. In contrast to HIV, where hypermutation was dominated by GG-to-AG/AA changes characteristic of A3G activity, the predominant hypermutation signature in SIV was GA-to-AA. Consistent with this shift, hypermutated SIV sequences contained substantially fewer premature stop codons than hypermutated HIV sequences. More importantly, even nonhypermutated sequences – the vast majority of viruses within the population – showed a clear enrichment of A3-associated mutations, particularly GA-to-AA relative to other mutation types. This enrichment was significantly greater in SIV than in HIV, demonstrating that A3 mutagenesis extends well beyond classical hypermutated genomes and contributes substantially to shaping the broader viral population.

Together, these results indicate that A3G deficiency in Cercopithecinae species reshapes the viral mutational landscape, shifting it from predominantly lethal hypermutation toward a mixture that includes sublethal mutagenesis. This shift promotes viral diversification and may contribute to differences in viral evolution, disease outcomes, and therapeutic responses between humans and NHPs.

Beyond A3G, the A3 gene family has diversified extensively in primates through domain duplication, gene fusion, retrocopying, and recurrent positive selection driven largely by retroviruses and retrotransposons.^93,94^ For example, NWMs encode up to ten *A3G* retrocopies, but lack *A3B*, *A3D*, and *A3F*.^95,96^ In OWMs, lineage-specific changes include altered subcellular localization of A3B, gained viral restriction activity of A3A in Cercopithecinae, a unique 27-amino-acid C-terminal tail and functional loss of A3H in primates, and differences in A3C activity between humans, great apes, and macaques.^97–101^ These variations underscore that A3-mediated antiviral immunity in NHPs diverges substantially from humans, with important translational implications given these enzymes roles in shaping the viral mutational landscape.

The implications of these findings extend beyond HIV and SIV. Notably, the reduction in A3G mutational signature GG-to-AG/AA has also been reported for foamy viruses in monkeys compared to those in zoonotically infected humans^53,54^ and also for simian T-cell leukemia virus type 1 (STLV-1) compared to human T-cell leukemia virus type 1 (HTLV-1),^52^ providing independent support for reduced A3G activity in these animal models. A3 enzymes are increasingly recognized as important drivers of mutation in a wide range of viral pathogens, including SARS-CoV-2, monkeypox virus, HPV, and EBV.^102–110^ Thus, understanding lineage-specific differences in A3 regulation is critical for accurately interpreting NHP infection models and for predicting host-virus interactions and viral evolutionary trajectories across diverse pathogens.

In summary, our study establishes the molecular basis, evolutionary origin, and functional impact of a lineage-specific *A3G* splicing defect in Cercopithecinae NHPs. By integrating comparative genomics, multiomic profiling, and functional analyses, we demonstrate that reduced A3G expression fundamentally reshapes viral hypermutation and evolution in these species relative to humans. Importantly, although many animals exhibited a complete A3G splicing defect, substantial inter-individual variation was observed, with some animals showing splicing defects as low as ∼40%. This variability suggests that animals with A3G expression profiles more closely resembling those of humans may exist within current NHP colonies. Our findings therefore highlight the importance of understanding and accounting for host genetic variation when selecting and interpreting these critical animal models.

Previous studies have emphasized the importance of genotyping NHPs and considering their substantial genetic diversity in biomedical research.^111–120^ Building on this foundation, we propose the concept of Precision Preclinical Medicine (PPM). This would involve leveraging natural genetic variation to identify individual animals whose biological characteristics closely match those of humans for the specific pathway, disease, or therapeutic question under investigation. This is analogous to clinical precision medicine, where molecular information guides patient care. PPM uses molecular information to guide model selection, with the goal of improving the reproducibility, translational fidelity, and biological relevance of preclinical research.

## Supporting information

S1-NHP-A3G-coordinates

S2-PCR-primers-spreading-infectivity-assay

S3-Total-spectral-counts-MSLdb-SwissProt-trypsin-2-LFDR-XT-Whole-PBMC-lysate

S4-Total-spectral-counts-MSLdb-SwissProt-LFDR-XT-gel-bands

## ACKNOWLEDGEMENTS

This research was supported by the Texas Developmental Center for AIDS Research (D-CFAR), a program funded by the National Institutes of Health (NIH) (5P30AI161943-03, D.E.); NIH/National Institute of Allergy and Infectious Diseases (NIAID) grants for HIV research (R01AI179465-01A1 and R56AI174877, D.E.); NIH/NIAID support for the HIV Accessory & Regulatory Complexes (HARC) Center (U54AI170792, J.F.H.); NIH support for the Third Coast Center for AIDS Research (P30AI117943, J.F.H.); additional NIH/NIAID grants for HIV research (R01AI167778, R01AI150455, R01AI165236, and R01AI150998, J.F.H.; R37-AI064046, R.S.H.); and the UT San Antonio Graduate School and NIAID training grant (T32AI184340, A.D.M.). R.S.H. is an investigator of the Howard Hughes Medical Institute, a CPRIT Scholar, and the Ewing Halsell President’s Council Distinguished Chair at the University of Texas at San Antonio Health Sciences Center. The following reagent was obtained through the NIH HIV Reagent Program, Division of AIDS, NIAID, NIH: Polyclonal Anti-Human APOBEC3G, C-terminal (antiserum, Rabbit), ARP-10201, contributed by Dr. Jaisri Lingappa and Simian Immunodeficiency Virus Type 1 (SIV-1) Infectious Molecular Clone, pSIVmac239 SpX, HRP-12249 contributed by Dr. Ronald C. Desrosiers. This project used biological materials funded by the National Center for Research Resources (P51 RR013986) and the base grant to the Southwest National Primate Research Center (P51 OD011133) from the Office of Research Infrastructure Programs, NIH. The content is solely the responsibility of the authors and does not necessarily represent the official views of the National Institutes of Health. We thank the Douglass Foundation Fellowship (Texas Biomedical Research Institute) for their generous support (A.D.M.). Mass spectrometry analyses were conducted with expert technical assistance of Sammy Pardo and Dana Molleur at UT San Antonio Institutional Mass Spectrometry Laboratory, under the direction of Susan T. Weintraub, Ph.D. The laboratory is supported in part by UTHSCSA, the University of Texas System Proteomics Core Network, which provided funds for the purchase of the Orbitrap Fusion Lumos mass spectrometer, and by NIH grant S10 OD030371-01A1 (S.T.W.), which supported the purchase of the Orbitrap Exploris 480 mass spectrometer. We are grateful to New Objective, Inc. for the custom-designed PicoFrit Resolve™ HPLC column used for analysis of the PBMC lysates. We also thank the University of Delaware DNA and Genotyping Center and Novogene for conducting PacBio sequencing. This work received computational support from Texas Biomed High Performance Computing Center (HPCC) and UT San Antonio’s HPCC, Arc, operated by Tech Solutions. The authors thank Dr. Tim Anderson for his insightful comments and constructive feedback on the manuscript, Sandy Smith, Gilbert Ramirez, and John Heaner for providing IT and computational support, Dr. Martha Lyke, Alec Bagwell, Dr. Ian Cheeseman, and Dr. Shelley Cole for providing rhesus macaque whole exome sequencing data under grant U42OD010442. Figures were created with BioRender.com.

## CONTRIBUTIONS

Conceptualization: A.D.M., R.S.-R., F.C.-T., J.F.H., D.E.; Data Curation: A.D.M., R.S.-R., A.B., F.C.-T., M.R., C.R., M.S., L.F.-P., S.T.W; Formal Analysis: A.D.M., A.B., F.C.-T., N.H., L.F.-P., M.S., S.T.W; Funding Acquisition: J.F.H., D.E.; Investigation: A.D.M., R.S.-R., F.C.-T., M.R., C.R., L.F.- P; Methodology: A.D.M., R.S.-R., A.B., F.C.-T., N.H., C.R., J.D.L., M.S., M.A.C., R.T.; Project Administration: R.S.-R., J.F.H., D.E; Resources: A.D.M., R.S.-R., F.C.-T., M.R., L.F.-P., H.Y., M.A.C., R.N.M., M.M., B.L., L.D.G., M.R.-B., R.S.H., X.C., J.F.H., D.E; Supervision: J.F.H., D.E; Validation: A.D.M., R.S.-R., F.C.-T., C.R., N.H.; Visualization: A.D.M., F.C.-T., N.H., J.D.L.; Writing - Original Draft Preparation: A.D.M., F.C.-T., J.F.H., D.E; Writing - Review & Editing: A.D.M., R.S.-R., A.B., F.C.-T., N.H., M.R., C.R., J.D.L., M.S., L.F.-P., H.Y., M.A.C., R.T., S.K., R.N.M., M.M., B.L., L.D.G., M.R.-B., R.S.H., X.C., S.T.W., J.F.H., D.E.

