## Supplemental_Materials for "APOBEC3G Splicing Defects in Nonhuman Primate Models Result in Disparate Viral Mutational Profiles Relative to Humans"

**SUPPLEMENTARY MATERIALS**

| *A3G* **Isoform** | **Full-length read counts** |
|---|---|
| PB.1708.5 | 5 |
| PB.1708.6 | 2 |
| PB.1708.4 ($A3G_{M2}$) | 20 |
| PB.1708.3 | 2 |
| PB.1708.1 | 9 |

**Supplementary Table S1**. The full-length read counts supporting each predicted A3G isoform generated from an Iso-Seq dataset of rhesus lymphoblastoid cell lines (*N* = 3) [63].

| Virus | Subtype | Molecule type | Sequences (*n*) | Considered (*n*) | Hypermutated (*n*) |
|---|---|---|---|---|---|
| HIV | All | All | 932,134 | 636,916 | 6,152 |
| HIV | All | DNA | 373,817 | 255,452 | 6,152 |
| HIV | All | RNA | 555,290 | 379,385 | 0 |
| SIV | All | All | 55,146 | 34,463 | 171 |
| SIV | All | DNA | 18,443 | 9,842 | 28 |
| SIV | All | RNA | 36,670 | 24,619 | 143 |
| SIV | MAC | All | 36,238 | 20,681 | 63 |
| SIV | MAC | DNA | 9,161 | 5,165 | 19 |
| SIV | MAC | RNA | 27,045 | 15,515 | 44 |
| SIV | SAB | All | 733 | 727 | 13 |
| SIV | SAB | DNA | 275 | 270 | 1 |
| SIV | SAB | RNA | 458 | 457 | 12 |
| SIV | SMM | All | 15,185 | 10,657 | 88 |
| SIV | SMM | DNA | 7,356 | 3,183 | 1 |
| SIV | SMM | RNA | 7,829 | 7,474 | 87 |
| SIV | TAN | All | 162 | 159 | 7 |
| SIV | TAN | DNA | 156 | 153 | 7 |
| SIV | TAN | RNA | 6 | 6 | 0 |

**Supplementary Table S2.** Descriptive characteristics of HIV and SIV sequences obtained from the Los Alamos National Laboratory (LANL) database.

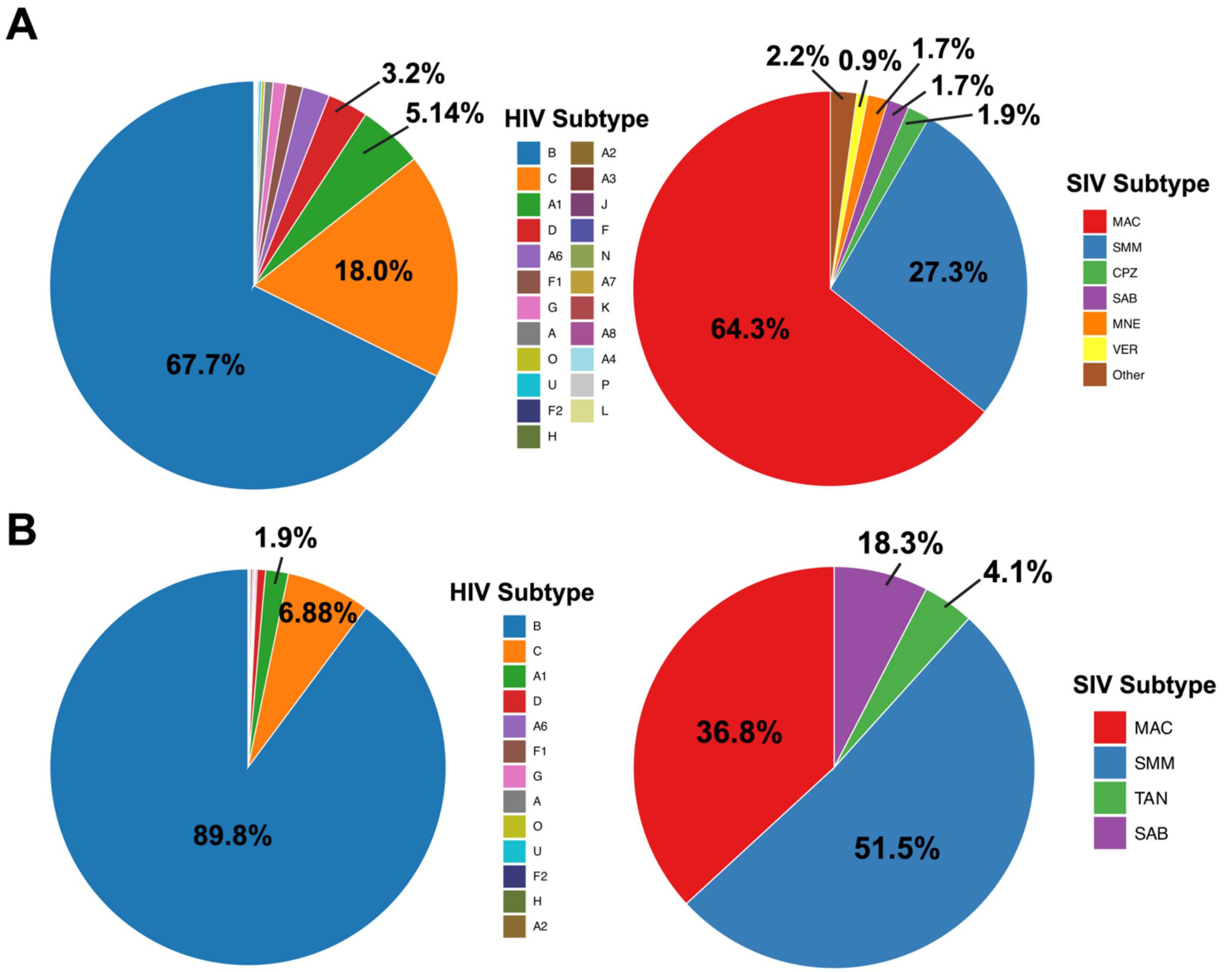


**Supplementary Figure S1.** Pie chart displaying the proportion of HIV and SIV subtypes **A)** All sequences **B)** Hypermutated sequences.

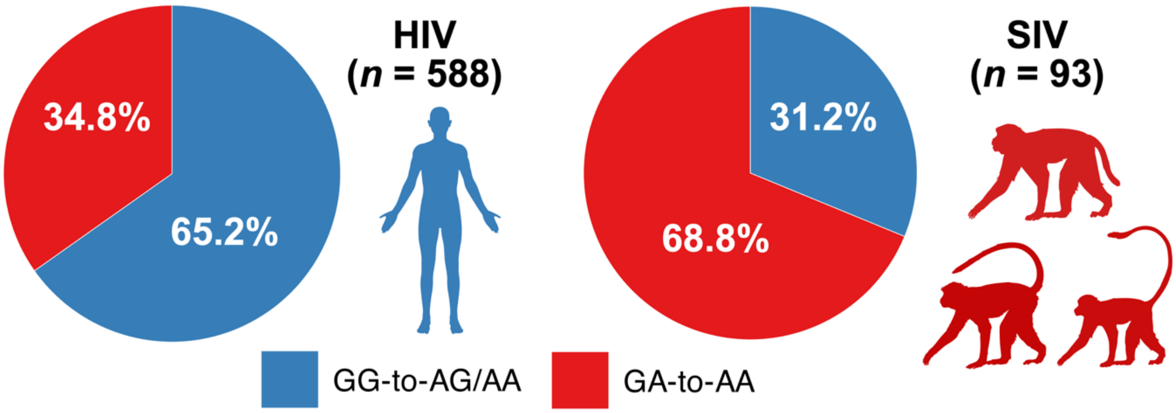


**Supplementary Figure S2. Divergent A3-associated hypermutation profiles in HIV and SIV**. Pie charts displaying the proportion of GG-to-AG/AA (%) and GA-to-AA (%) mutations averaged across all hypermutated sequences within each donor.

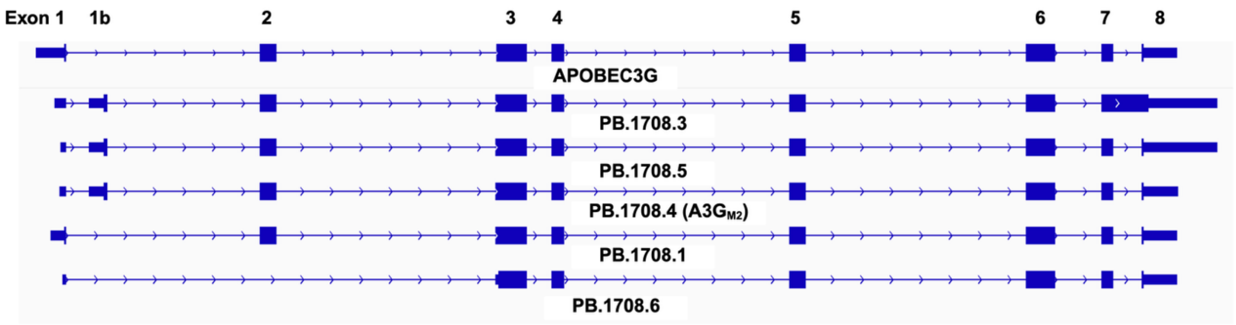


**Supplementary Figure S3**. **Multiple *A3G* isoforms identified using PacBio IsoSeq**. Long-read defined transcript models for rhesus macaque A3G identified in lymphoblastoid cell lines (LCLs) (*N* = 3) [63].

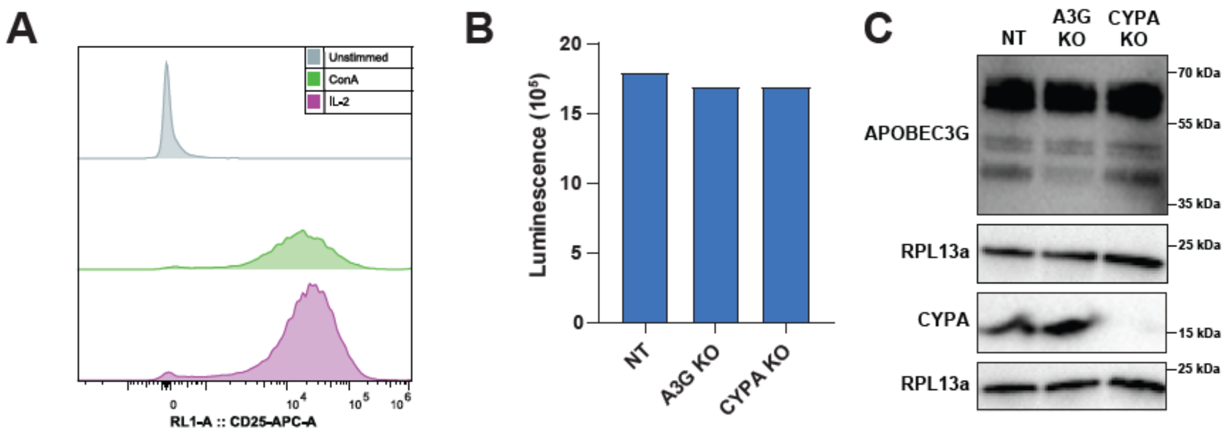


**Supplementary Figure S4. Knock-out of A3G in rhesus macaque PBMCs. A)** Validation of CD4+ T cell stimulation after Con A treatment by anti-CD25 immunostaining and flow cytometry. **B)** Cell viability after electroporation with crRNPs by a luminesce based ATP assay (CellTiterGlo). **C)** Immunoblots for rhesus macaque A3G, CYPA, and RPL13a (loading control). Results from 1 female animal shown here.

| Peptide | Primate | Abundance ratio |
|---|---|---|
| IYDDQGR | Human | 1.040 |
| IYDDQGR | Chimpanzee | 0.362 |
| IYDDQGR | Gorilla | 0.885 |
| IYDDQGR | Orangutan | 1.914 |
| IYDDQGR | Rhesus macaque | 0.147 |

**Supplementary Table S3**. **Among the primate studied, rhesus has the lowest A3G-specific peptide abundance ratio**. A3G peptide abundance ratios for human and various NHPs revealed by mass spectrometry.[63]

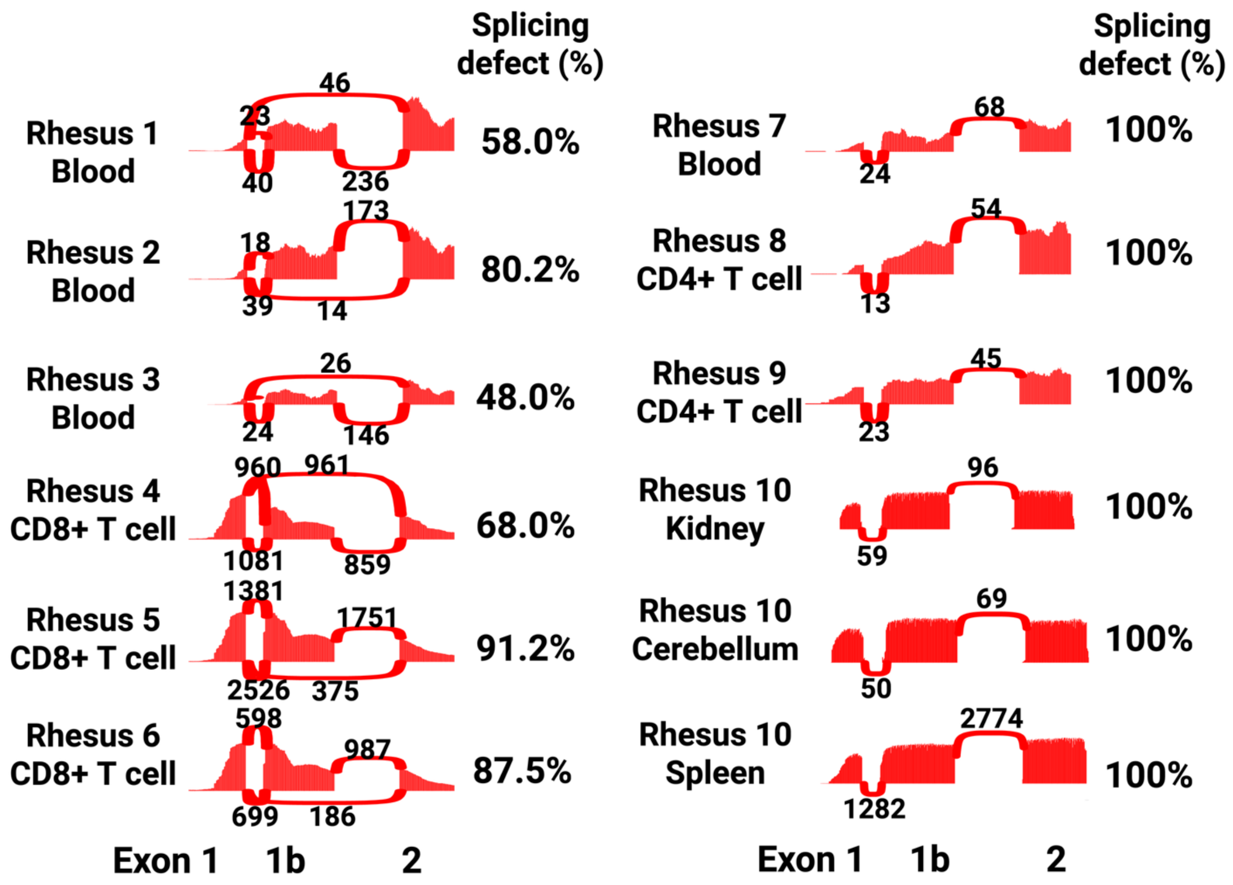


**Supplementary Figure S5. Aberrant *A3G* mRNA splicing in rhesus macaque is very common**. A3G Sashimi plots from diverse rhesus macaques across different studies indicate that inclusion of exon 1 in *A3G* mRNA is widespread. Splicing defect (%) is calculated by (# of reads that map from exon 1 to exon 1b / # of reads that map from exon 1 to exon 1b + # of reads that map from exon 1 to 2) * 100.

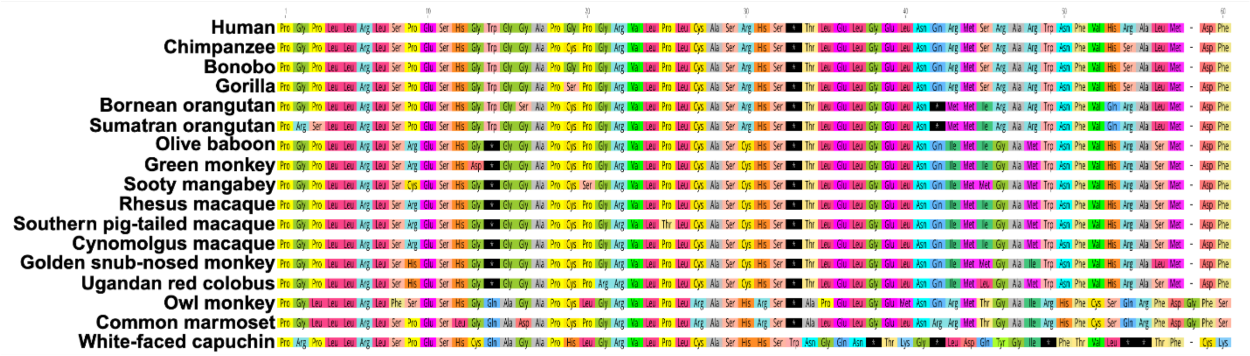


**Supplementary Figure S6. Two premature termination codons (PTCs) are present in A3G exon 1b across all studied Cercopithecinae subfamily members**. Exon 1b protein alignment (using MUSCLE) showing that two stop codons (represented by black*) are present in all Cercopithecinae subfamily members.

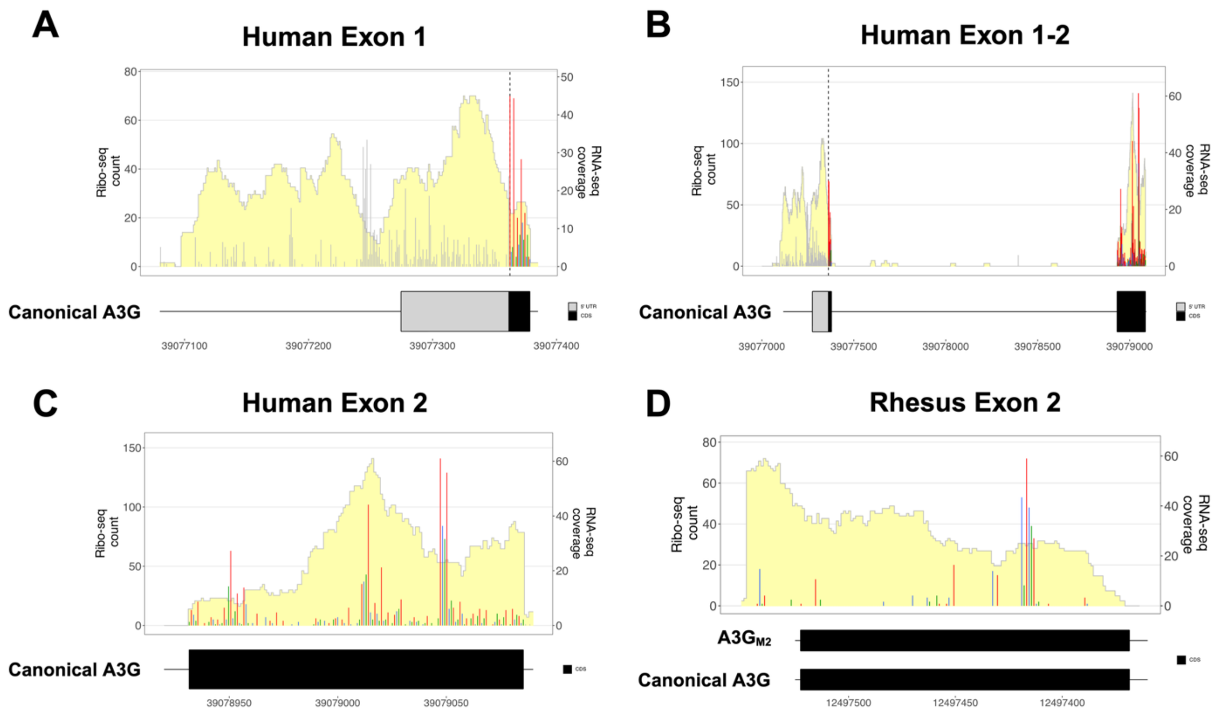


**Supplementary Figure S7. Human A3G displays active translation as indicated by codon periodicity.** Ribo-seq and matched RNA-seq coverage of **A)** human *A3G* exon 1, **B)** human *A3G* exons 1-2, **C)** human *A3G* exon 2, and **D)** rhesus macaque *A3G* exon 2. RNA-seq coverage is shaded in yellow and Ribo-seq counts are determined by P-site calculation and are displayed as grey (not in the main annotated coding sequence (CDS)), red (frame 0), blue (frame 1), or green (frame 2) peaks.

| Species | Gene ID | RPKM | TPM |
|---|---|---|---|
| **Rhesus macaque** | ENSMMUG00000039346 | 13.5125 | 19.0167 |
| **Human** | ENSG00000239713 | 36.2781 | 42.8785 |

**Supplementary Table S4. Human A3G has more ribosome footprints than rhesus macaque in A3G transcripts from Ribo-seq data.** Normalized Ribo-seq counts of *A3G* in rhesus macaque ($N$ = 5) and human ($N$ = 4).

| **Gene ID** | **Transcript ID** | **Genome Position (chr:start-end:strand)** | **Ribo *p*-value** | **Frame *q*-value** | **Amino acid length** |
|---|---|---|---|---|---|
| ENSMMUG00000039346 (*A3G*) | A3G$_{M1}$ | 10:12489051-12499017:- | 2.145e-07 | 4.88e-06 | 394 |
| ENSMMUG00000039346 (*A3G*) | A3G$_{M2}$ | 10:12489051-12499029:- | 2.145e-07 | 4.88e-06 | 394 (390) |

**Supplementary Table S5. Ribo-TISH open reading frame (ORF) prediction for rhesus A3G$_{M1}$ and A3G$_{M2}$ isoforms.** Despite different CDS annotations in the gene transfer format (GTF) file, Ribo-TISH assigns A3G$_{M2}$ a 394–amino-acid ORF because it overlaps the A3G$_{M1}$ ORF.

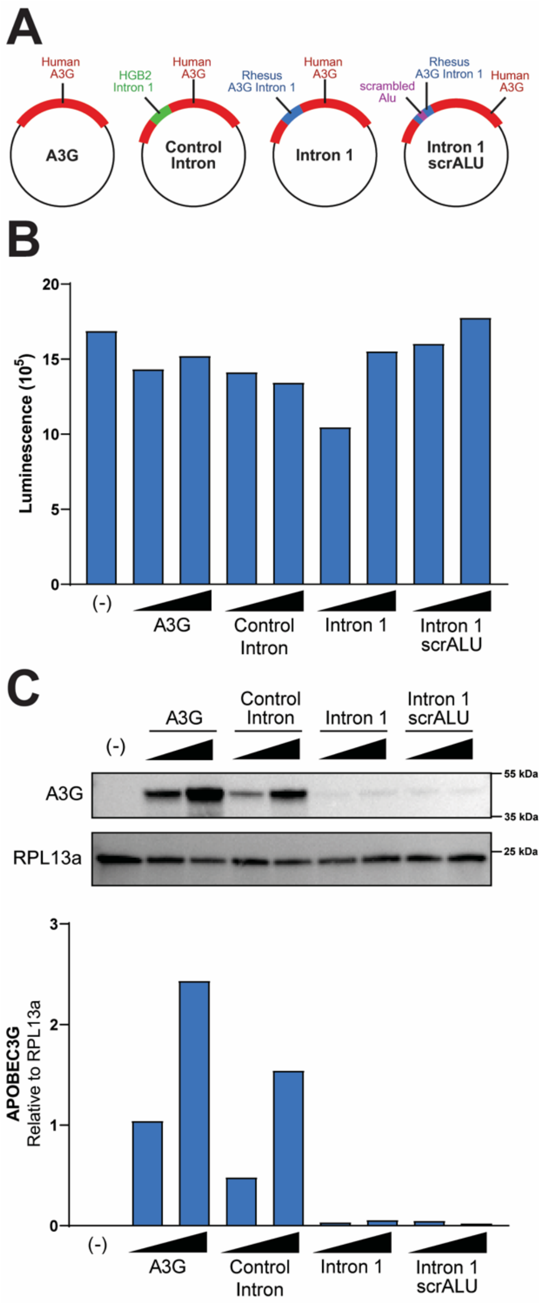


**Supplementary Figure S8. Scrambling the *Alu* sequence does not rescue A3G protein expression. A)** Schematic of plasmid constructs, **B)** Cell viability of HEK293T cells expressing constructs assessed by CellTiter-Glo. **C)** Immunoblots of HEK293T cells expressing indicated plasmid constructs, and quantification relative to loading control RPL13a, showing that rhesus intron 1 strongly ablates A3G expression and that scrambling the *Alu* sequence does not rescue A3G protein expression.

| Virus | Donor | Molecule type | PCR | Sequences mapped (*n*) | Hypermutated (*n*) | Proportion (%) |
|---|---|---|---|---|---|---|
| HIV | Hu3 | ALL | ALL | 1387250 | 18236 | 1.31 |
| HIV | Hu3 | DNA | ALL | 1212034 | 17727 | 1.46 |
| HIV | Hu3 | DNA | NFL | 559597 | 7926 | 1.42 |
| HIV | Hu3 | DNA | env | 450336 | 7234 | 1.61 |
| HIV | Hu3 | DNA | gag | 202101 | 2567 | 1.27 |
| HIV | Hu3 | RNA | ALL | 175216 | 509 | 0.29 |
| HIV | Hu3 | RNA | env | 67938 | 398 | 0.59 |
| HIV | Hu3 | RNA | gag | 107278 | 111 | 0.10 |
| SIV | ALL | ALL | ALL | 3897637 | 44796 | 1.15 |
| SIV | ALL | DNA | ALL | 1797608 | 37837 | 2.10 |
| SIV | ALL | DNA | NFL | 996420 | 24996 | 2.51 |
| SIV | ALL | DNA | env | 534375 | 10743 | 2.01 |
| SIV | ALL | DNA | gag | 266813 | 2098 | 0.79 |
| SIV | ALL | RNA | ALL | 2100029 | 6959 | 0.33 |
| SIV | ALL | RNA | env | 1254828 | 6350 | 0.51 |
| SIV | ALL | RNA | gag | 845201 | 609 | 0.07 |
| SIV | Rm5 | ALL | ALL | 1534505 | 21774 | 1.42 |
| SIV | Rm5 | DNA | ALL | 857165 | 19215 | 2.24 |
| SIV | Rm5 | DNA | NFL | 500588 | 12832 | 2.56 |
| SIV | Rm5 | DNA | env | 238037 | 5269 | 2.21 |
| SIV | Rm5 | DNA | gag | 118540 | 1114 | 0.94 |
| SIV | Rm5 | RNA | ALL | 677340 | 2559 | 0.38 |
| SIV | Rm5 | RNA | env | 262050 | 2111 | 0.81 |
| SIV | Rm5 | RNA | gag | 415290 | 448 | 0.11 |
| SIV | Rm7 | ALL | ALL | 2363132 | 23022 | 0.97 |
| SIV | Rm7 | DNA | ALL | 940443 | 18622 | 1.98 |
| SIV | Rm7 | DNA | NFL | 495832 | 12164 | 2.45 |
| SIV | Rm7 | DNA | env | 296338 | 5474 | 1.85 |
| SIV | Rm7 | DNA | gag | 148273 | 984 | 0.66 |
| SIV | Rm7 | RNA | ALL | 1422689 | 4400 | 0.31 |
| SIV | Rm7 | RNA | env | 992778 | 4239 | 0.43 |
| SIV | Rm7 | RNA | gag | 429911 | 161 | 0.04 |

**Supplementary Table S6.** Descriptive characteristics of HIV and SIV sequences by donor, molecule type, PCR region, the number of sequences mapped, and the number of hypermutated sequences identified from the spreading infectivity assay. NFL = near-full-length

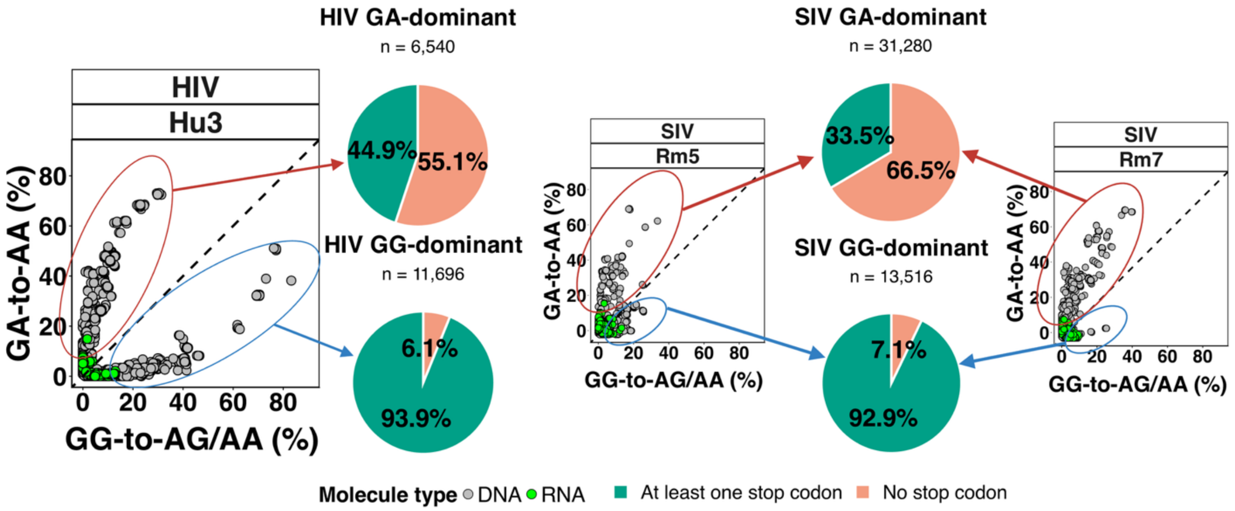


**Supplementary Figure S9. GG-dominant hypermutated sequences are much more likely to be defective than GA-dominant hypermutated sequences.** The proportion of hypermutated sequences with at least one stop codon segregated by GG-dominant or GA-dominant sequences in HIV and SIV. GG-dominant sequences (circled in blue) are defined as sequences in which the burden of GG-to-GA>AA is greater than the burden of GA-to-AA. GA-dominant sequences (circled in red) are defined as sequences in which the burden of GA-to-AA is greater than the burden of GG-to-AG.
